# The interplay of uncertainty, relevance and learning influences auditory categorization

**DOI:** 10.1101/2022.12.01.518777

**Authors:** Janaki Sheth, Jared S. Collina, Eugenio Piasini, Konrad P. Kording, Yale E. Cohen, Maria N. Geffen

## Abstract

Auditory perception requires categorizing sound sequences, such as speech or music, into classes, such as syllables or notes. Auditory categorization depends not only on the acoustic waveform, but also on variability and uncertainty in how the listener perceives the sound – including sensory and stimulus uncertainty, the listener’s estimated relevance of the particular sound to the task, and their ability to learn the past statistics of the acoustic environment. Whereas these factors have been studied in isolation, whether and how these factors *interact* to shape categorization remains unknown. Here, we measured human participants’ performance on a multi-tone categorization task and modeled each participant’s behavior using a Bayesian framework. Task-relevant tones contributed more to category choice than task-irrelevant tones, confirming that participants combined information about sensory features with task relevance. Conversely, participants’ poor estimates of task-relevant tones or high-sensory uncertainty adversely impacted category choice. Learning the statistics of sound category over both short and long timescales also affected decisions, biasing the decisions toward the overrepresented category. The magnitude of this effect correlated inversely with participants’ relevance estimates. Our results demonstrate that individual participants idiosyncratically weigh sensory uncertainty, task relevance, and statistics over both short and long timescales, providing a novel understanding of and a computational framework for how sensory decisions are made under several simultaneous behavioral demands.

## Introduction

Making sensory decisions in everyday settings is a complex task due to the presence of multiple forms of uncertainty^1–8^. One of many such decisions is identifying what a friend said when the conversation takes place in a crowded restaurant versus in a quiet room. Despite the apparent ease by which we can accomplish this task, it is a very complicated computational process involving many factors. First, the sensory transduction process and neural-processing stages are noisy, generating a form of uncertainty called *sensory uncertainty*^9–12^. A second form of uncertainty is *stimulus relevance*: of all stimuli within a sensory scene, only some are relevant to the current decision^13–18^. For example, when trying to identify a particular sound uttered by our friend, we have to disregard sounds uttered by other speakers, which can be considered to be distractors. A third form of uncertainty is *stimulus uncertainty*, which is a listener’s uncertainty in estimating the variability in a sound source’s generation of the nominal same stimulus^19^. Even once our sensory system isolates the signals that were produced by our friend from background noise or irrelevant signals, we may be uncertain as to what specific sound or word was produced. Furthermore, when presented with any form of uncertainty, observers may rely on *the learned past statistics of stimuli*^20–26^ to make their decisions, such as their long-term knowledge of how their friend typically talks or their short-term knowledge of their friend’s recent speech sounds^27^. Indeed, observers can learn both *short-*^28,29^ and *long-term stimulus statistics*^30–32^ and this learning is considered crucial for efficient decision making^33,34^, with or without a conscious effort from the listener.

Whereas the effects of the factors – sensory uncertainty, stimulus relevance, stimulus uncertainty and short- and long-term statistics – on sensory decision-making, such as categorization, have previously been studied *separately*^19,35^, in everyday situations we often confront them *simultaneously*. Thus, these factors may *interact* with each other, differentially affecting category decisions. For example, sensory uncertainty may not only decrease our ability to categorize a sound uttered by our friend but may also affect our estimate of the relevant versus the distractor sounds. Our prior expectations of what our friend is saying may play a greater role when our sensory uncertainty or stimulus uncertainty is high or when there are more distracting sounds in the environment by other speakers. These expectations may be reduced if we are talking to a person we just met.

To test whether and how the interplay of these factors shapes sensory decisions, we measured the performance of human participants on an auditory categorization task^36–39^. More specifically, our goals were to test how observers’ sensory uncertainty and associated estimates of the likelihood of the irrelevant stimuli mediated their sensitivity to stimulus relevance, and how their task performance was affected by learning. Participants categorized trial sequences of three tones as either high frequency or low frequency in a two-alternative forced choice task, as we varied the number of task-irrelevant (distractor) tones and task-relevant (category-specific) tones across the trials. Additionally, to test learning, in a subset of experimental sessions, we modulated the proportion of trials biased toward the high or the low category.

Because it has been suggested that perceptual systems optimally integrate information from relevant inputs while ignoring the irrelevant ones^40^, we characterized participants’ behavior using normative Bayesian models of sensory decision-making^41–46^. The Bayesian formalism is a well-established framework to understand complex interactions among many factors governing decision-making^13,14,30,31,47^. On average, when categorizing auditory stimuli, participants integrated information across all three tones but weighed them by their task relevance. Stimulus uncertainty was similar across participants. However, individual participants’ sensitivity to task relevance depended on their internal sensory uncertainty and subjective estimates of the likelihood of irrelevant stimuli. We also found population-wide evidence of both learning short-term statistics (i.e., stimulus category of the previous trial) and learning long-term statistics (i.e., session-level bias toward high versus low category). However, individual participants’ knowledge of short- and long-term statistics was inversely correlated with their estimates of the likelihood of irrelevant stimuli. Together, our results demonstrate that humans differentially combine information about sensory uncertainty, stimulus relevance, stimulus uncertainty, and short-term and long-term learning, providing an integrated framework for understanding how the interplay of different mechanisms shapes auditory processing.

## Results

### Participants weigh tones according to stimulus relevance

Auditory categorization can depend on multiple factors, such as sensory uncertainty, stimulus relevance, stimulus uncertainty, and stimulus statistics that can vary over both short and long timescales. Because these factors often co-occur, they may interact with each other to affect decision-making and considering their relations they, indeed, computationally should interact. Here, we designed a categorization task to measure how these factors interacted in decision-making. In a two-alternative forced-choice auditory task, participants were asked to report the category of a three-tone sequence (Fig 1A). Each trial was randomly selected to be from low or high category, and each tone of the trial could probabilistically either be a signal or distractor (Fig 1B-D).

**Fig. 1:**
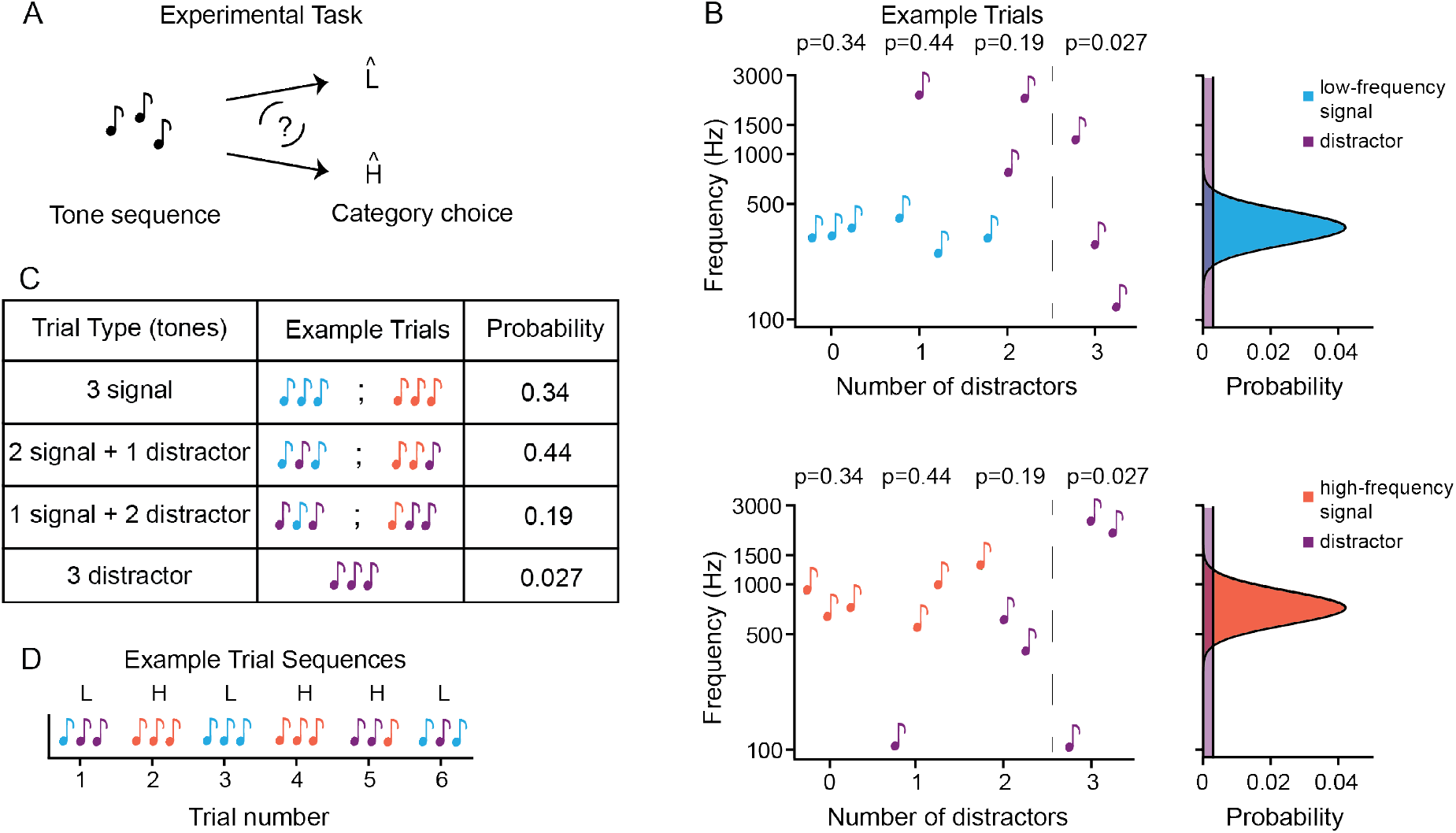
Schematic diagram of the auditory categorization task. (A) A participant categorizes a given three-tone trial sequence as high or low. (B) Example low (top) and high (bottom) category trials from an *unbiased* session. The underlying probability distribution for signal tones is Gaussian (low-frequency Gaussian: light blue; high-frequency Gaussian: red) and for distractor tones is uniform (purple). The stimuli in each trial are three tones denoted by notes: signal tones from the low-frequency distribution (light blue), signal tones from the high-frequency distribution (red), and distractor tones from the uniform distribution (purple). (C) The 4 types of trial combinations in the task and their corresponding probabilities. (D) Example trial sequences from one of the participant’s data in the *unbiased* session.

We first tested whether participants accounted for relevance of individual tones when categorizing the tone sequences. Because the total number of tones was the same during each trial, when a signal tone was replaced by a distractor tone, it decreased the amount of available information to the listener regarding stimulus category and added irrelevant information. To account for this, we analyzed trials separately with the same number of distractors across individuals (Fig. 2A,B). Compared to signal tones, distractors should have less of an impact on participants’ decisions, because they have no relevance for the task. As expected, participant’s category choice probability was more strongly correlated with the frequency of the presented signal tones compared to the frequency of the distractor tones (Fig 2A; two-sided paired t-test, trials with one distractor: *p* = 1.32e-22; trials with two distractors: *p* = 5.36e-18; Figs S1C-H and Table 1). This confirms that participants used tone relevance when making decisions.

**Table 1:**
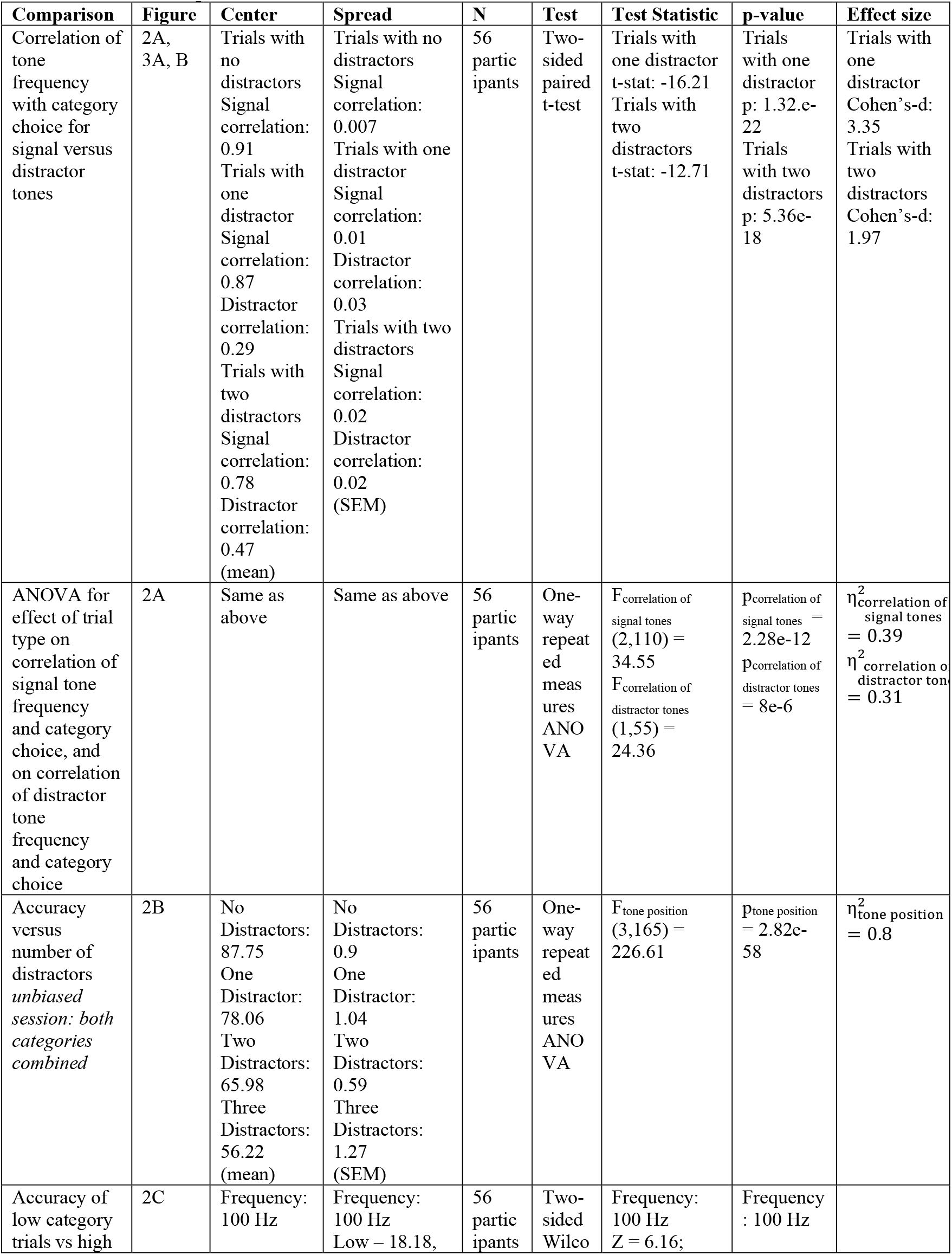
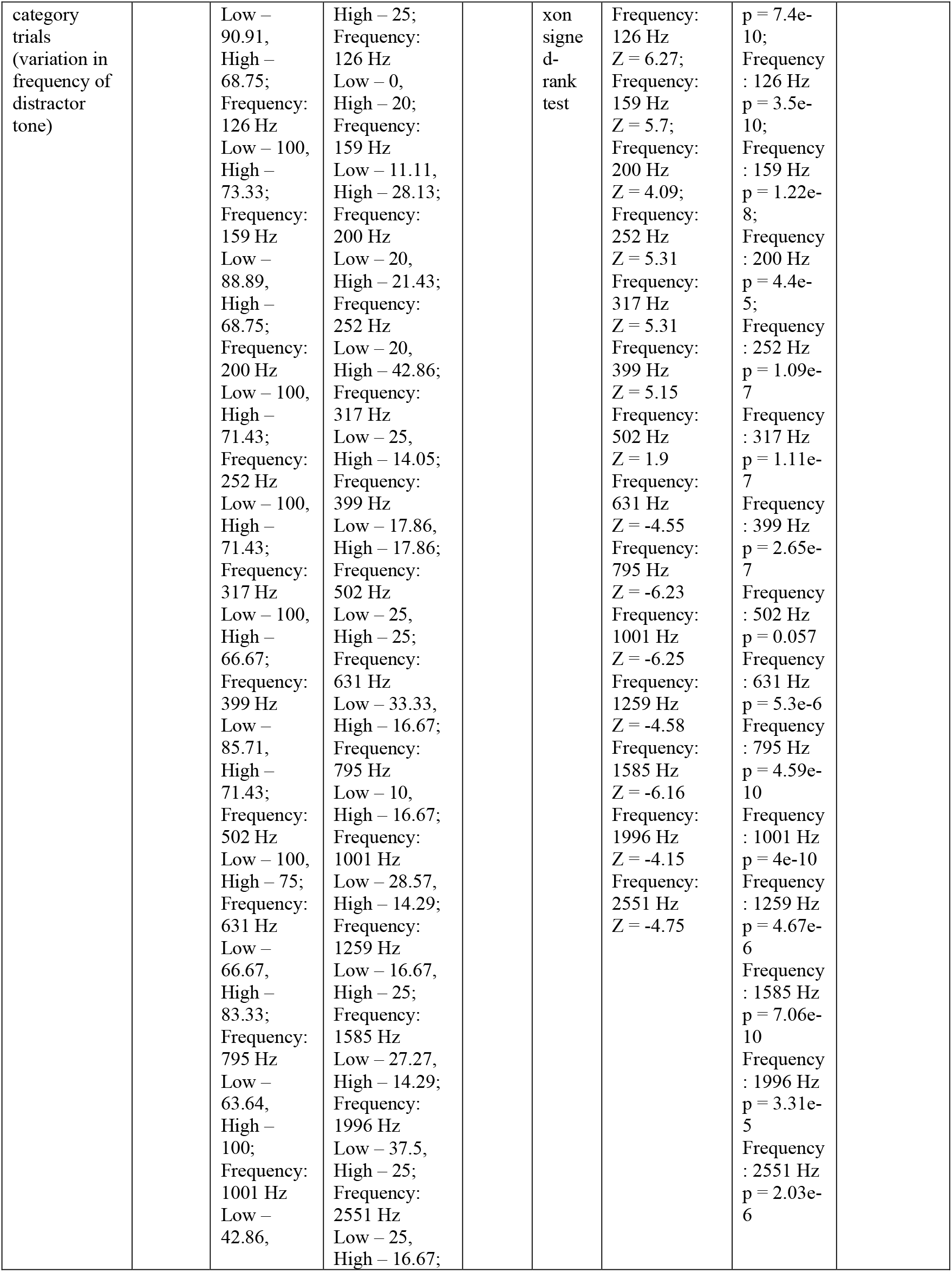
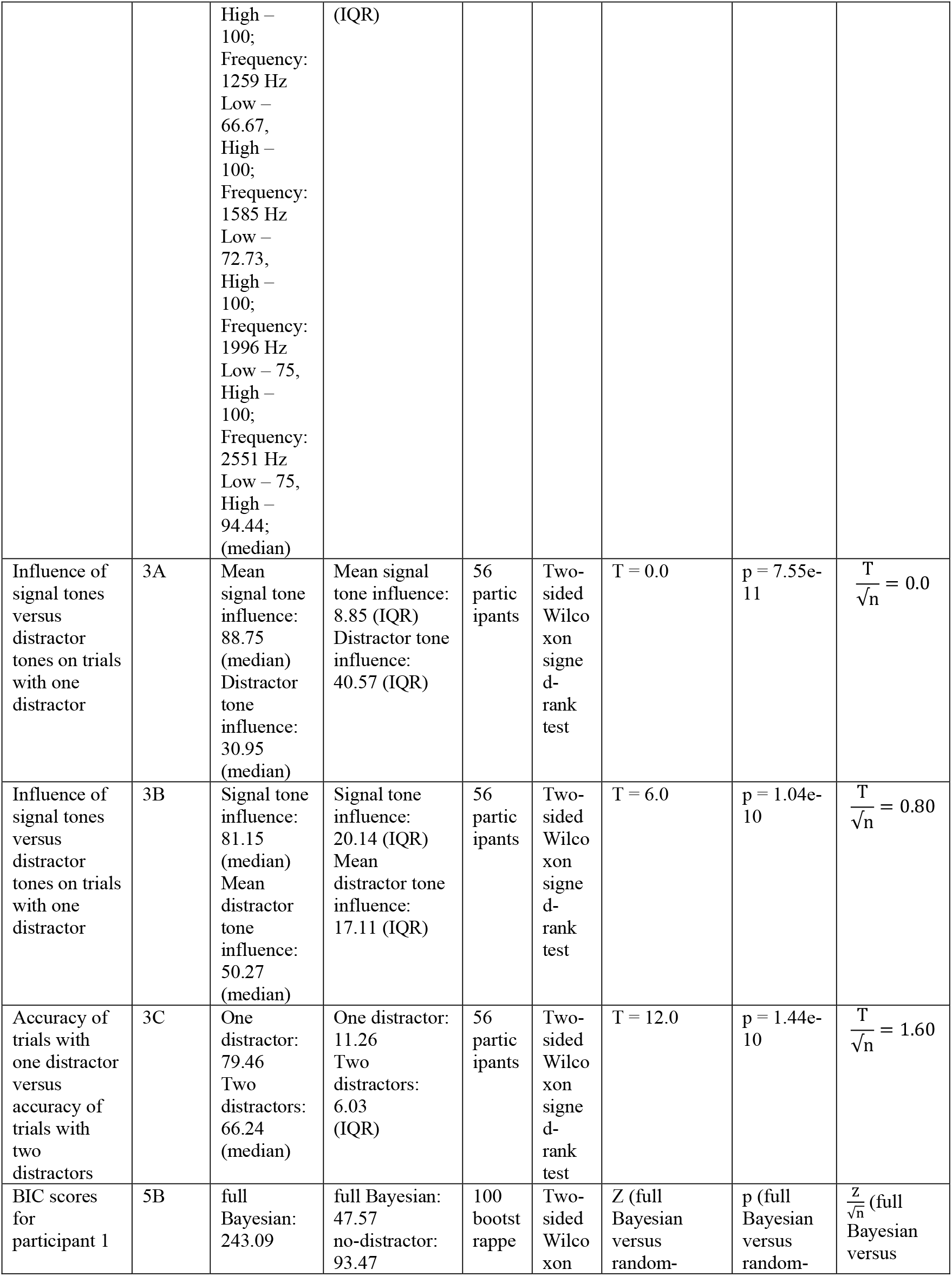
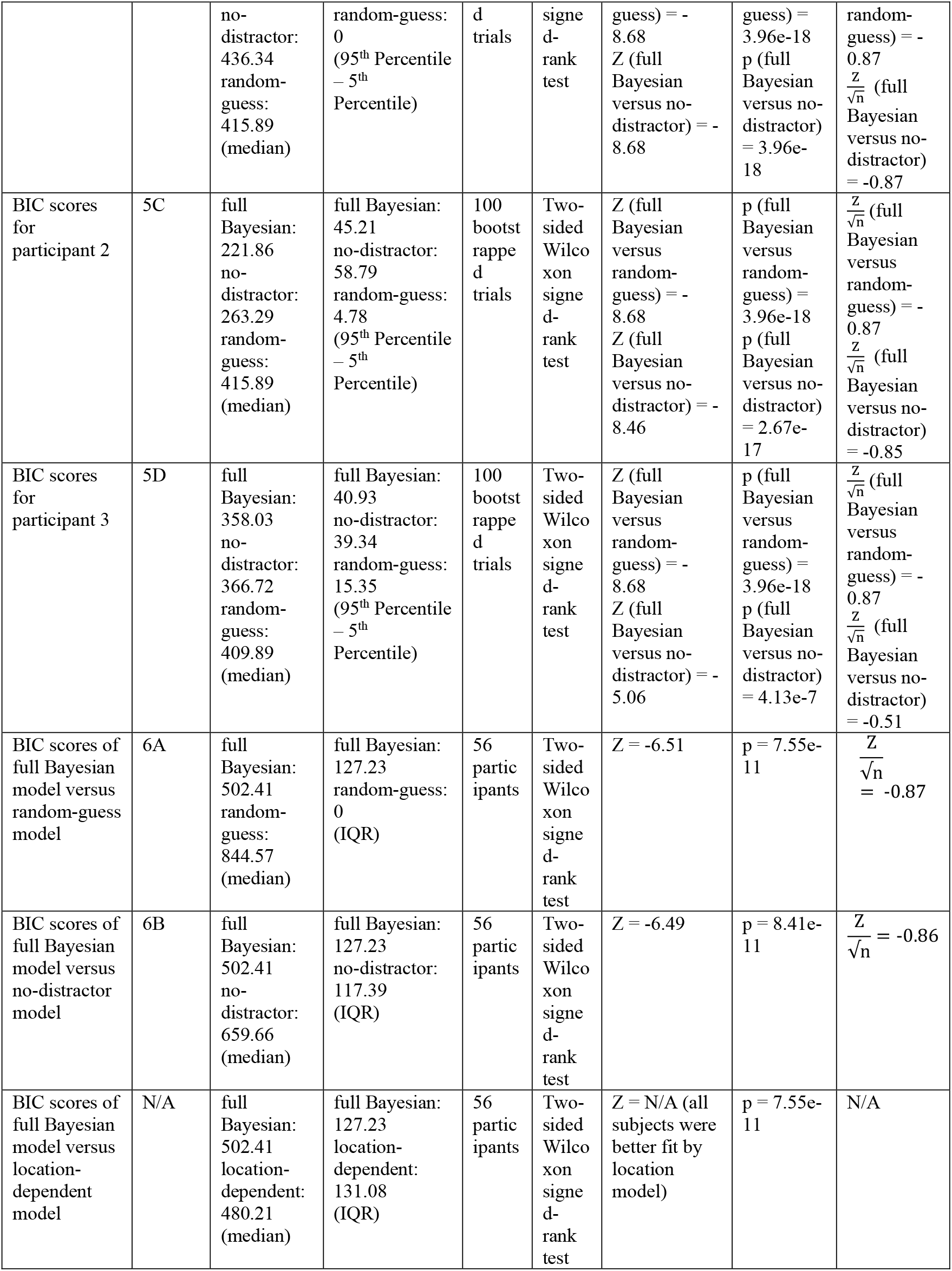
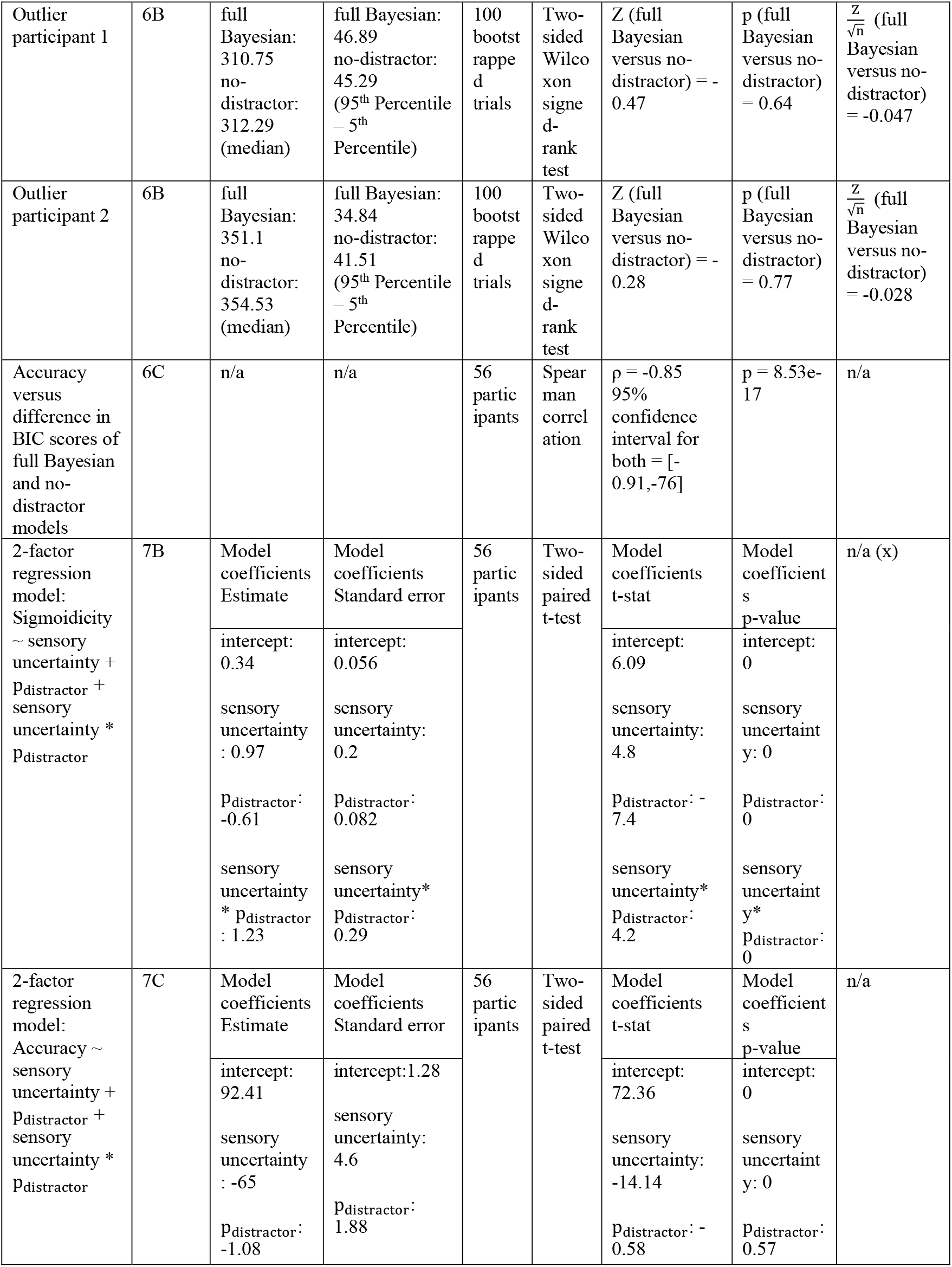
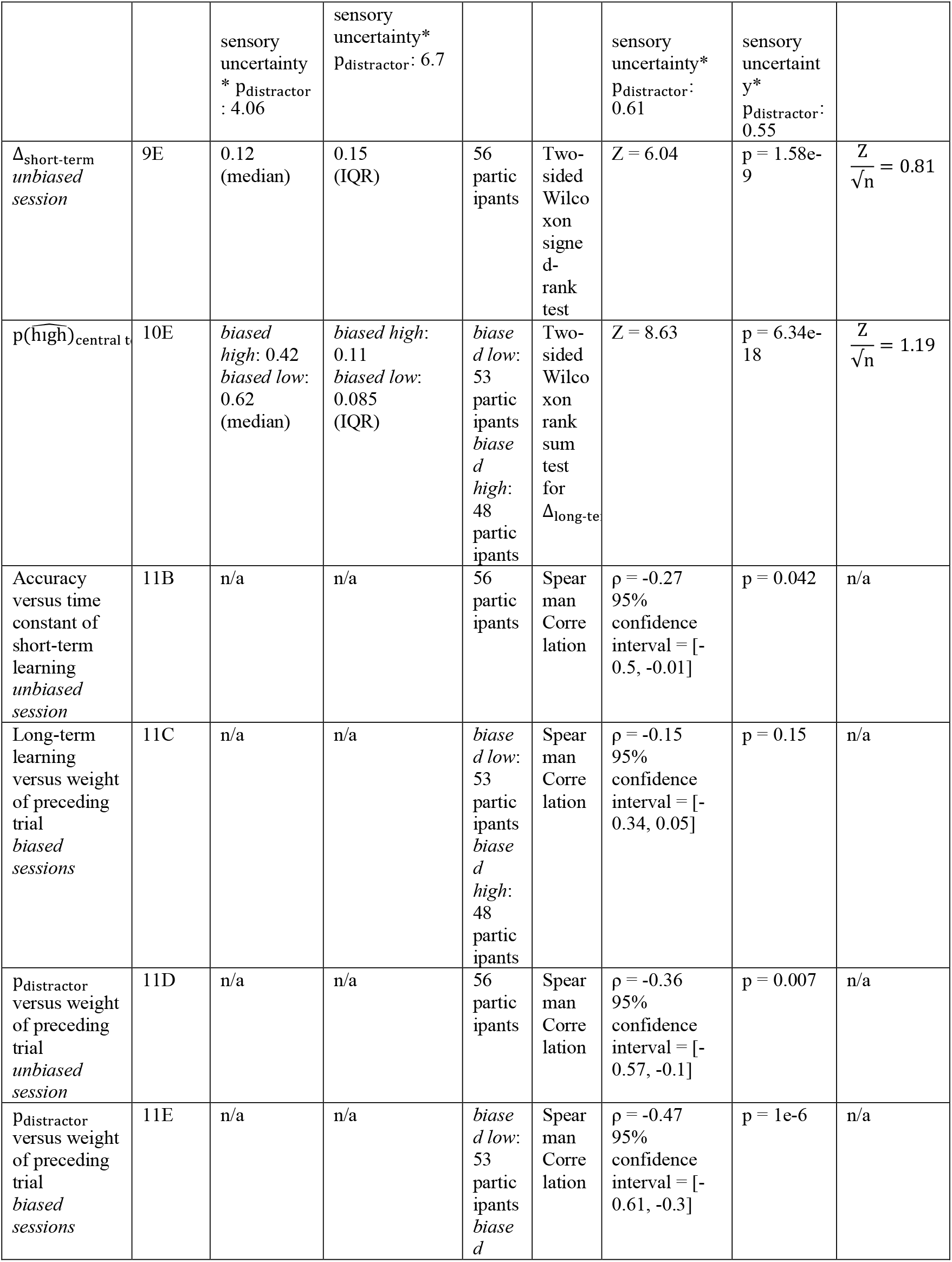
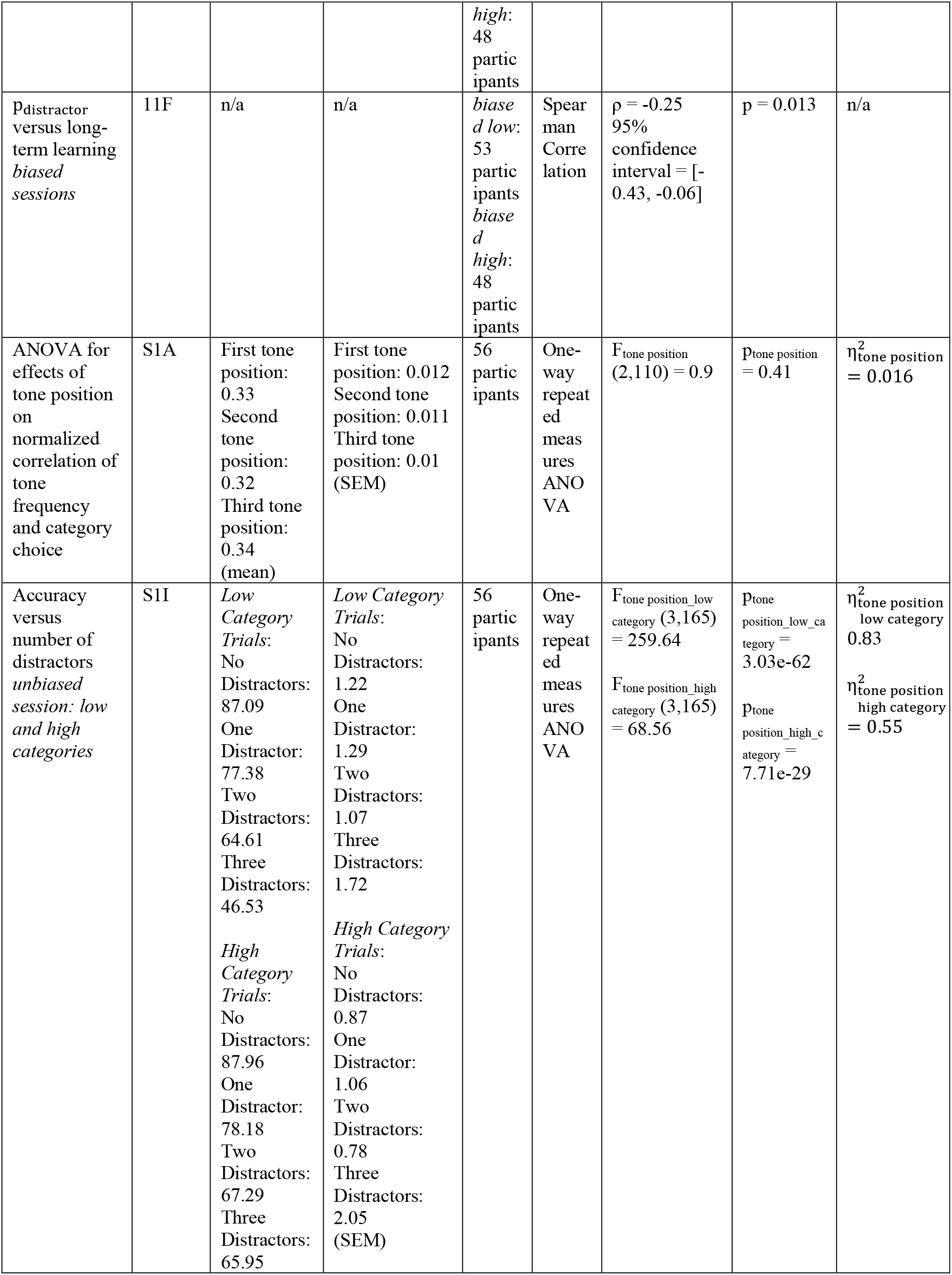
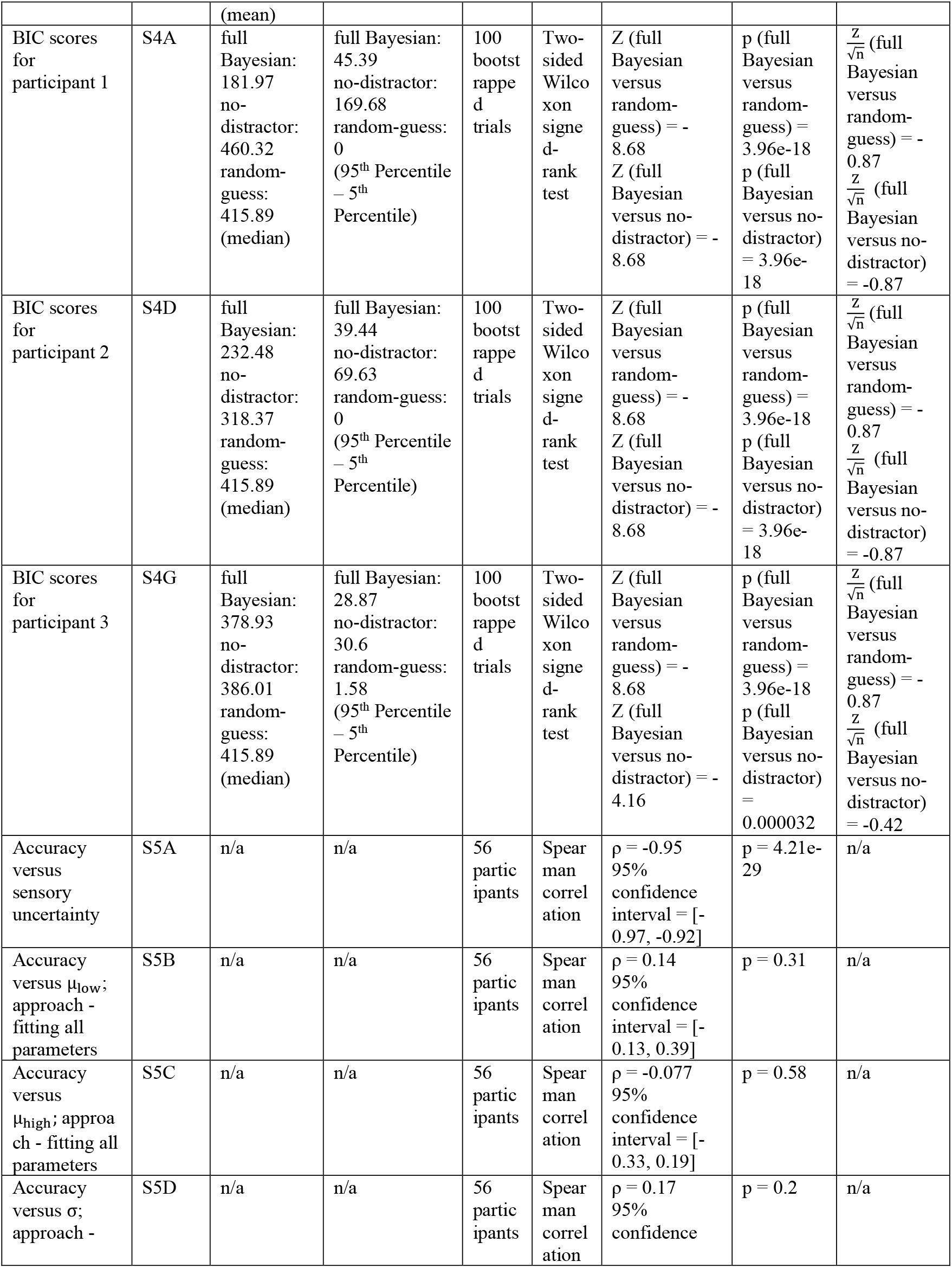
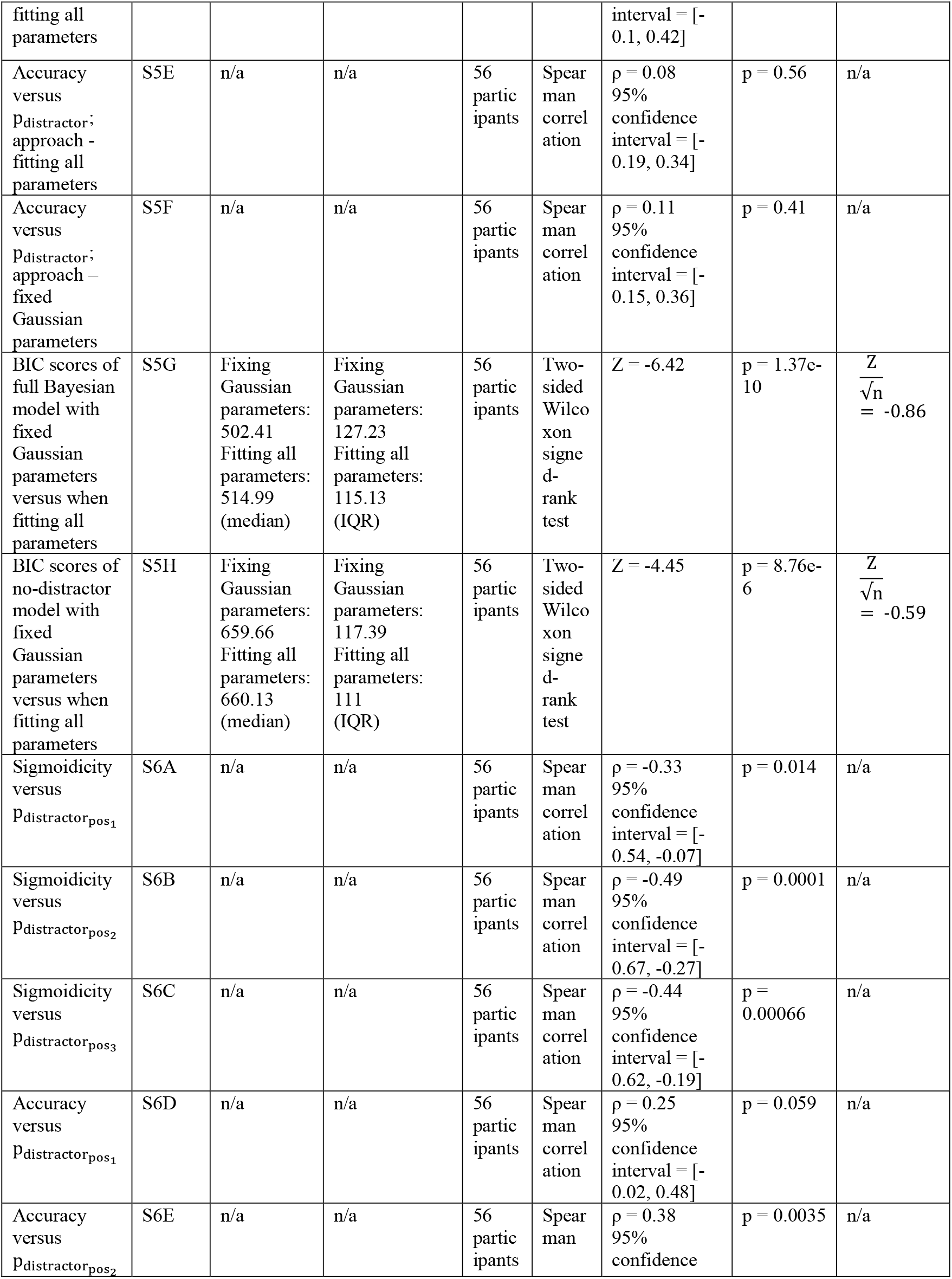
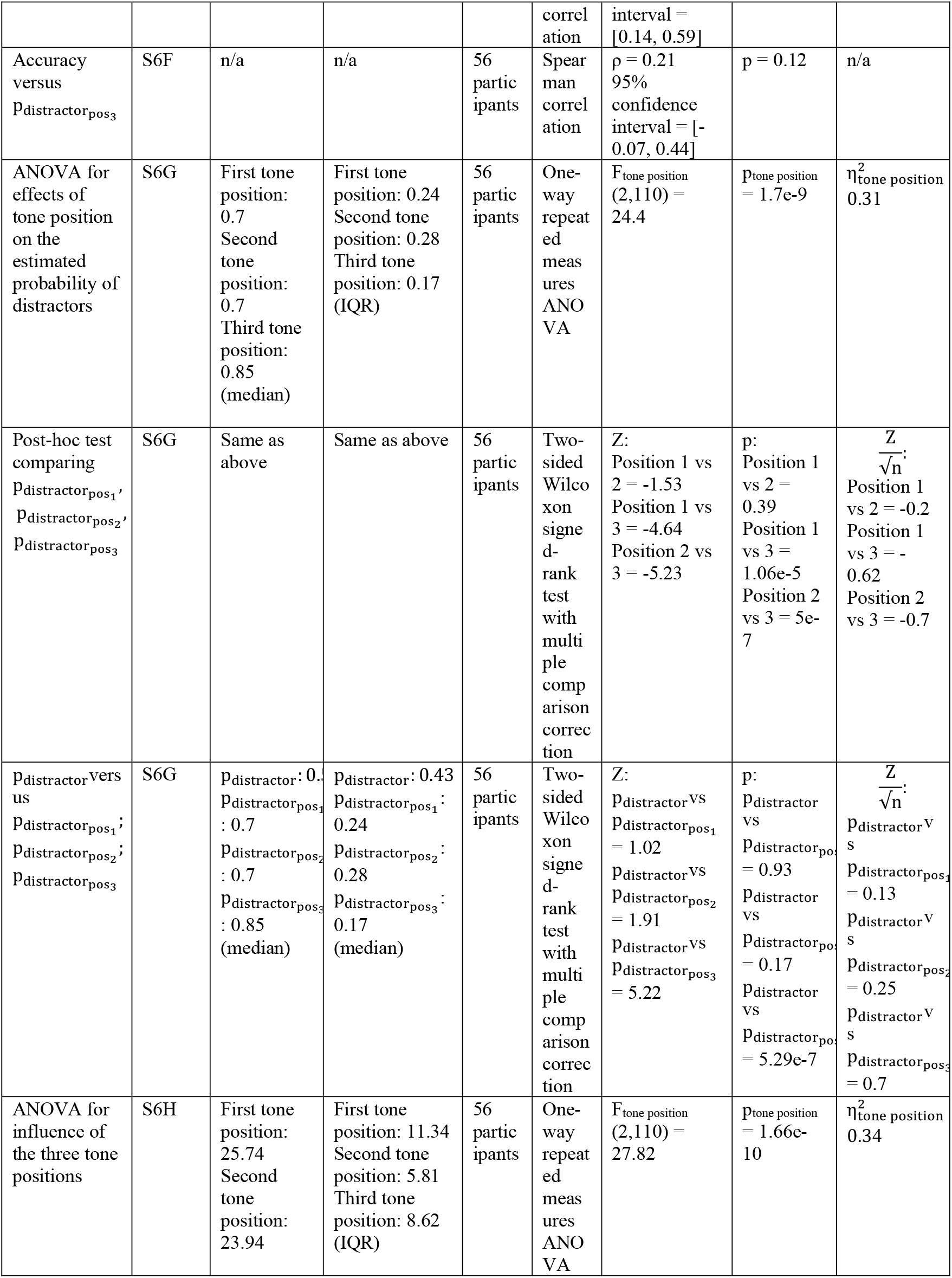
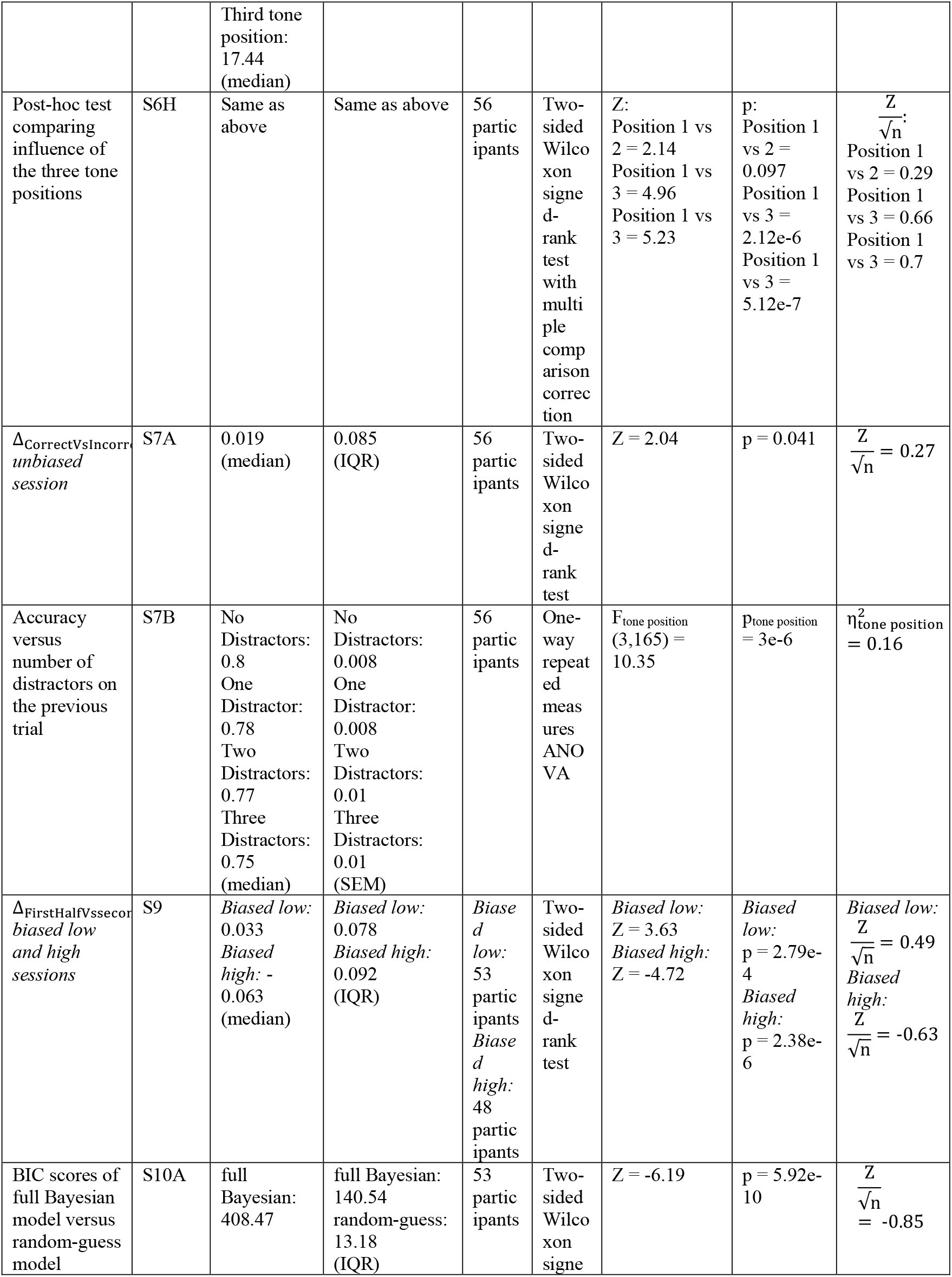
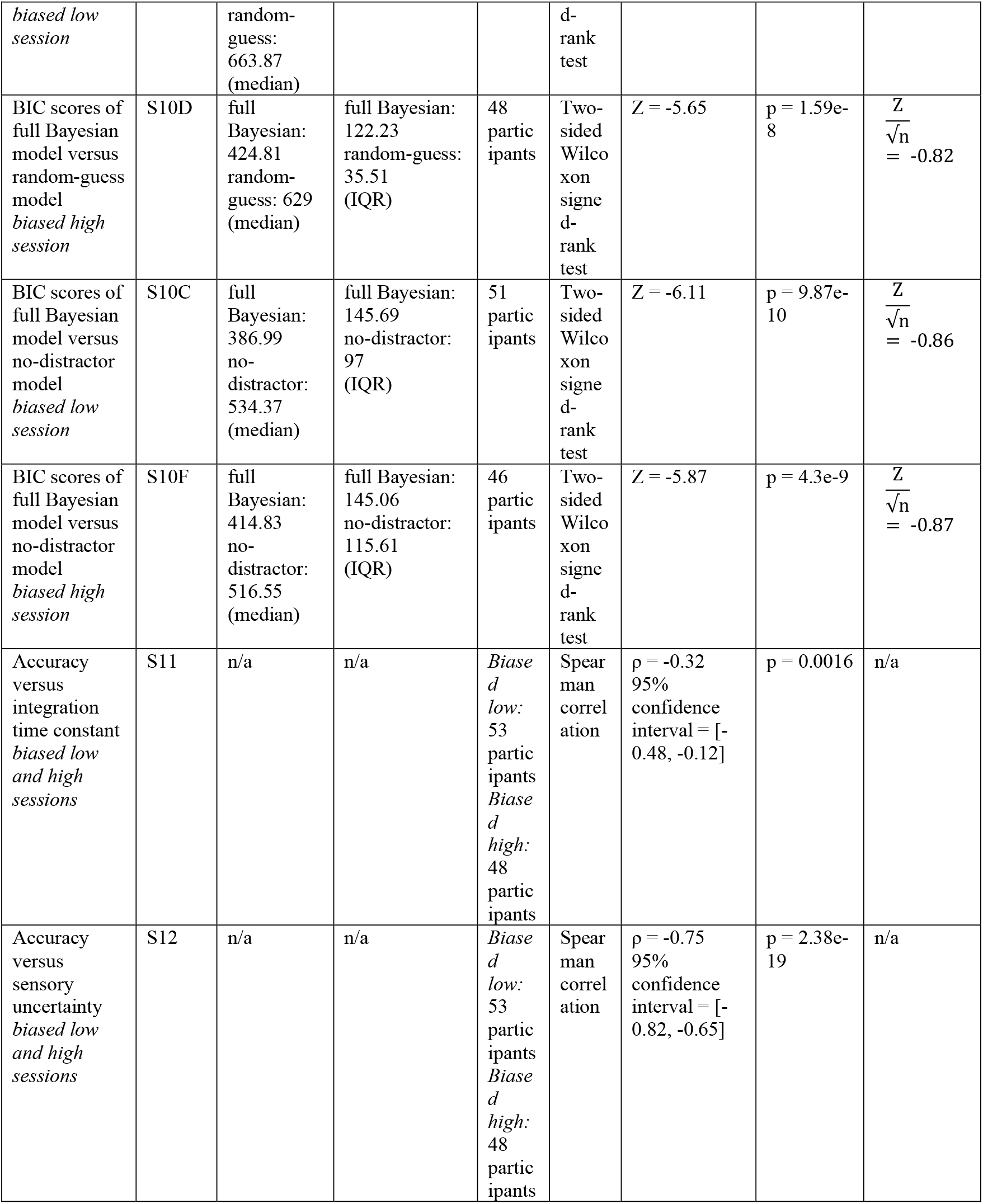
Statistical comparisons.

**Fig. 2:**
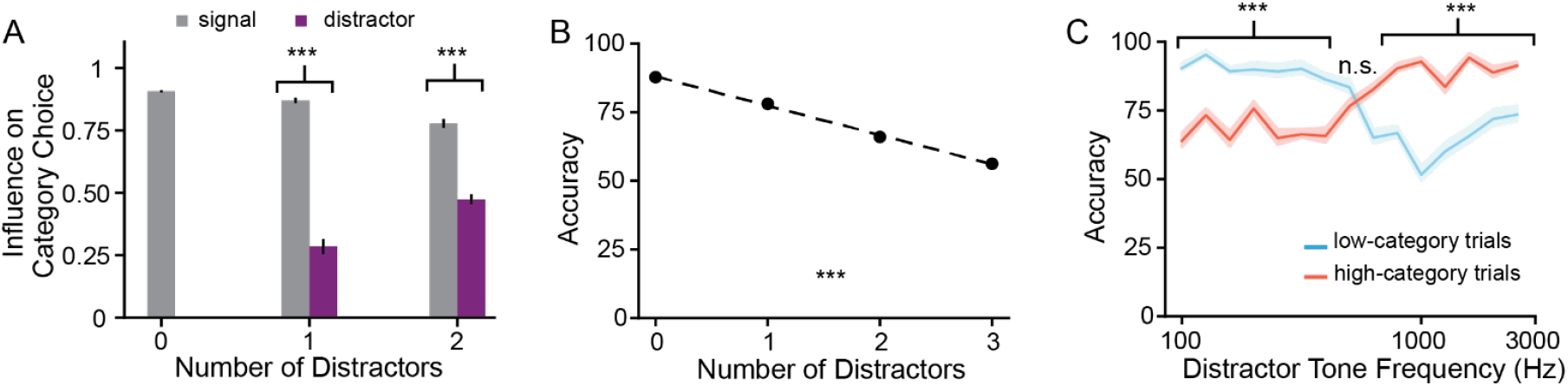
On average, participants weighted tones according to their relevance when making category decisions. (A) For both signal (grey) and distractor (purple) tones, their influence on category choice is computed using correlation of the respective tone frequencies with their associated average category choice probability (Methods). Error bars: standard error of the mean. Statistics: two-sided paired t-test between each subjects’ normalized correlation coefficient of signal tone frequencies with choice probability and correlation coefficient of distractor tone frequencies with choice probability. (B) Average accuracy in *unbiased* trials decreased with the number of distractor tones. Error bars: standard error of the mean. Statistics: one-way repeated measures ANOVA; p_tone_position_ = 2.82e-58, F_tone_position_(3, 165) = 226.61. (C) For trials with exactly one distractor tone, accuracy was higher when the distractor tone frequency was similar to the trial category (Blue: low-category trials, red: high-category trials). Binned frequencies of the distractor are shown on the x-axis. Error bars: standard error of the mean. Statistics: two-tailed Wilcoxon signed-rank test between accuracy in high- and low-category trials computed at each distractor frequency. N = 56 participants. ^†^p<0.05, *p<0.01, **p<0.001, ***p<0.0001.

Conversely, because distractor tones may still be integrated in the decision with some probability^48,49^, they may impair participants’ performance by providing information that should have been ignored. Indeed, as the number of distractors increased, the correlation between distractor tones and category choice, which we interpret as the influence of distractor tones on category choice, also increased whereas that of signal tones decreased (Fig 2A; Figs S1B-D; one-way rmANOVA, significant main effect for number of distractors on correlations with signal tone frequencies: *F(2,110)* = 34.55, *p* = 2.28e-12, and on correlations with distractor tone frequencies: *F(1,55)* = 24.36, *p* = 8e-6). Additionally, accuracy decreased as the number of distractors increased (Fig 2B; N = 56, correlation of accuracy with the number of distractors: Spearman, ρ = -1, *p* = 0).

These findings collectively demonstrate that because participants were uncertain about the relevance of the different tones, the distractors adversely impacted their decisions.

Next, we tested whether the effect of distractors depended on their tone frequency. We hypothesized that a distractor tone whose frequency was in the same frequency range as the trial’s generative category center would impair participants’ decisions less than a distractor tone whose frequency was not in the same frequency range. We evaluated this prediction by computing the average participant accuracy for trials with one distractor tone. Category accuracy for these trials was significantly higher when the frequency of the distractor was similar to the trial category (Fig 2C; Table 1 – Wilcoxon signed-rank test). For trials in which the distractor frequency was different than the trial category, we plotted separate psychometric curves corresponding to the signal or distractor tone frequencies (Fig S1J). A qualitative comparison of these curves to the psychometric curve for trials with only signal tones (Fig S1B) indicates that distractors have a substantial effect on participants’ category choice. Combined, these data suggest that participants’ ability to differentiate between signal and distractor tones depends on the frequency value of the distractors.

### Participants exhibit high individual variability in performance

Whereas the previous analysis confirmed the expected effects of tone relevance on the mean performance of the participants, the extent to which tone relevance impacts choice varies idiosyncratically across individuals. This diversity may be due to participants’ distinct internal priors, estimates of relevance, measure of stimulus uncertainty or their sensory uncertainty. Indeed, there was significant heterogeneity in the correlation of tone frequency with category choice probability (for both signal and distractor tones, Figs 3A, B) and with accuracy (across trials with one and two distractors, Fig 3C). Accuracy for trials with one distractor tone ranged from 57% - 89% and for trials with two distractor tones ranged from 53% - 75%. Thus, in subsequent analyses, we not only quantified the effects of relevance, stimulus uncertainty, sensory uncertainty, and learning of stimulus statistics on the average performance across the population but also on inter-participant variability. This allowed us to probe how these different real-life factors might mediate individual participant’s decision-making process.

**Fig. 3:**
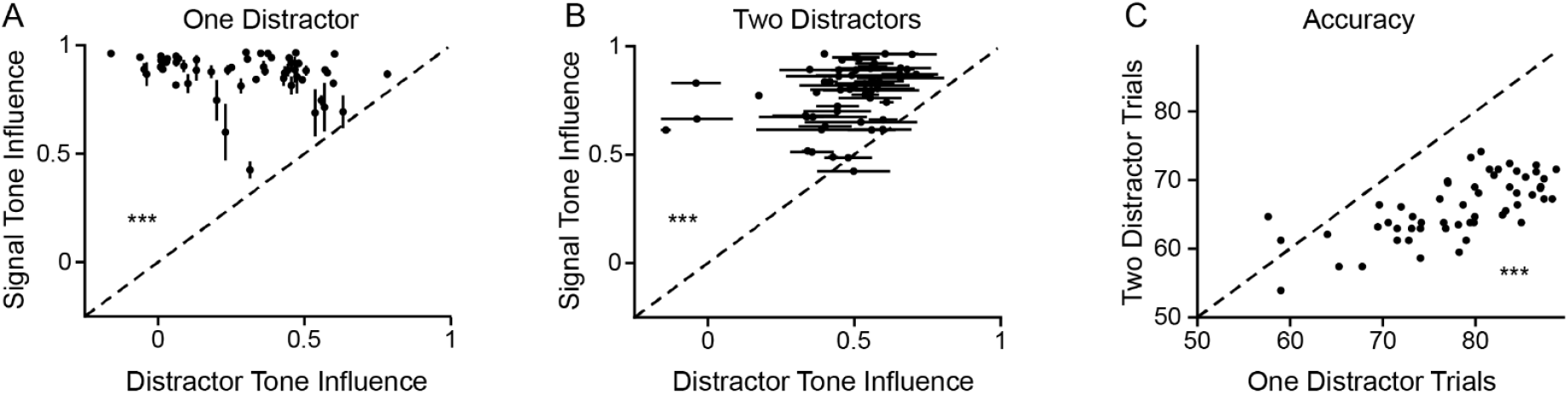
Individual participants differed in their estimate of relevance. (A, B) For each participant, influence of tone type on category choice is computed using the correlation of respective tone frequencies with category choice probability. (A) Influence of signal versus distractor tones for trials with one distractor. Statistics: two-tailed Wilcoxon signed-rank test for the 56 participants. (B) Influence of signal versus distractor tones for trials with two distractors. Statistics: two-tailed Wilcoxon signed-rank test for the 56 participants. (C) Categorization accuracy for the tone sequences with one versus two distractor tones. Statistics: two-tailed Wilcoxon signed-rank test for the 56 participants. Error bars: standard error of the mean obtained by assuming that a participant’s accuracy follows a binomial distribution; not large enough to be distinguishable. ^†^p<0.05, *p<0.01, **p<0.001, ***p<0.0001.

### Bayesian model can account for participant performance

So far, we have demonstrated in a model-free way that when making decisions, participants across the population accounted for the relevance of the different tone frequencies as they integrated information from the three tones. Next, to investigate how these factors shaped individual participant’s category decisions, we modeled each participant’s behavior using a Bayesian approach. We first discuss how tone frequency should affect individual category choice probability and then lay out three models for understanding decisions and their predictions for this relation.

On a standard two alternative forced choice (2AFC) frequency categorization task with no distractor tones, tone frequencies at the intersection of the two gaussian distributions do not provide evidence for either category. As the tone frequencies move further away from the decision boundary, tones become more informative about the trial’s category (Figs 4A, B) and consequently, a participant’s accuracy increases the further the tone frequencies are from the decision boundary. Thus, in the absence of distractors, if a trial consisted of a single tone, a participant’s choice probability would be modeled by a standard psychometric curve, which is a monotonically increasing saturating sigmoid. In contrast, if the trial consisted of multiple tones, choice probability could be modeled as the product of the relevant sigmoidal single-tone psychometric curves.

**Fig. 4:**
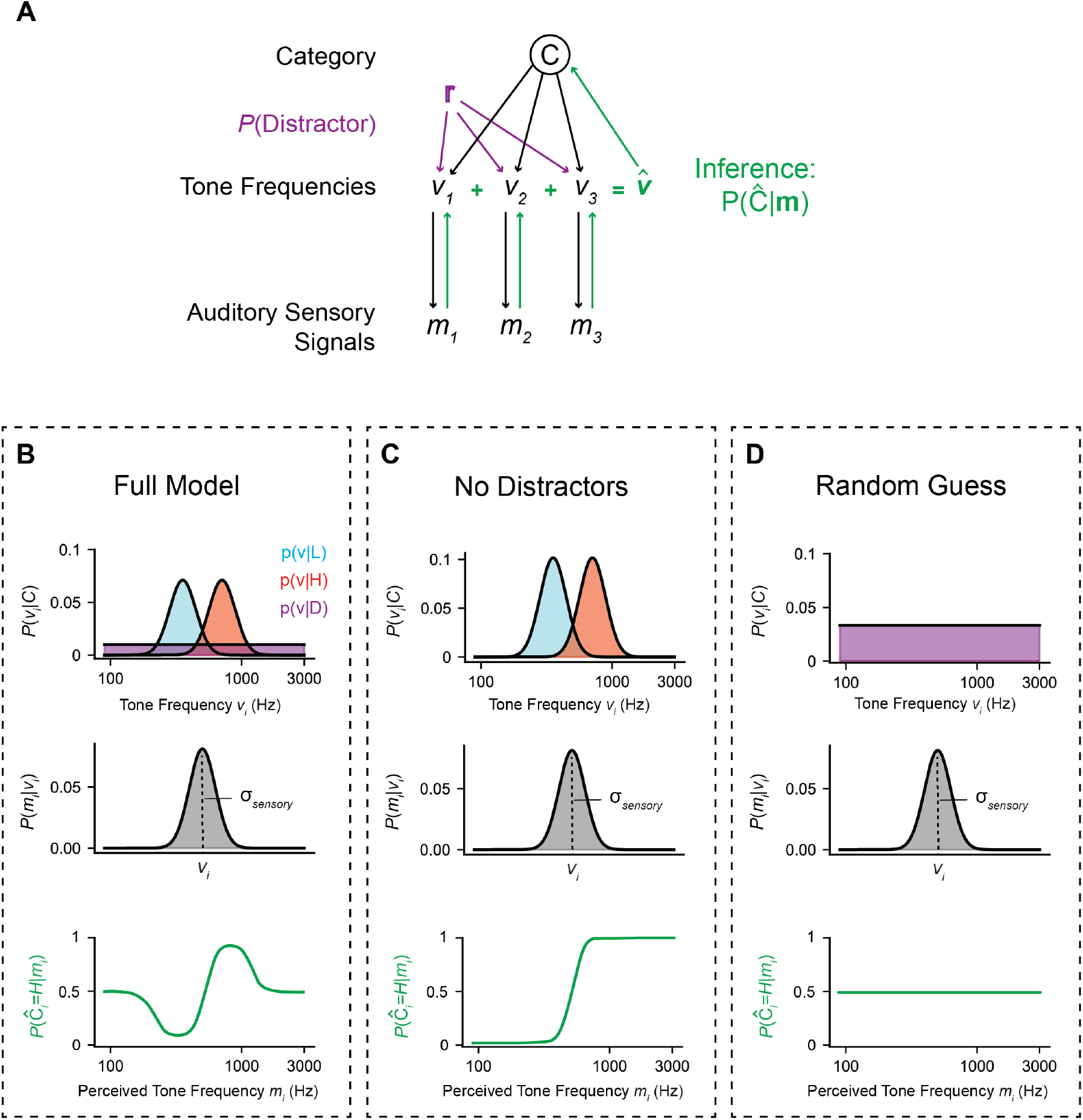
Graph of the Bayesian Model. (A) The (top level) category identity C of the tone-burst sequence constrains the values of the (middle level) three individually generated tone frequencies v_i_. Each tone frequency has a probability of 30% of being replaced by a distractor frequency uniformly drawn from the full frequency range. The (bottom level) auditory sensory signal m_i_ represents a noisy measurement of the true tone frequency v_i_. The black arrows define the generative conditional probability densities p(v|C) and p(m|**v**). The task of the observer is to infer the category membership of the full tone sequence **v** from the noisy sensory measurement **m** = {m_1_, m_2_, m_3_} while considering the possibility of distractor tone bursts (green arrows). (B) The Full Bayesian Model considers participants to be veridical about the parameters of the generative distribution and the distractor probability. Top: Given a particular category, the perceived probability of a certain tone frequency is governed by the respective conditional distribution p(v|C) and the probability of that tone being replaced by a distractor tone p_distractor_. Middle: The sensory process of the Bayesian observer is modeled as a Gaussian process centered at the true stimulus frequency. Bottom: If the tones can be judged to be either signal or distractor, the psychometric curve becomes ‘S’-shaped. Further, this curve plateaus at 0.5 because frequency values at the tails of the Gaussian distributions are more likely to be distractors than signal tones. (C) The No Distractors Model considers those participants that believe that there are no distractors, and therefore all perceived stimuli are drawn from either the “high” or “low” distributions. Top: Given a particular category, the perceived probability of a certain tone frequency is governed by the respective conditional distribution p(v|C). Middle: The sensory process of the Bayesian observer is modeled as a Gaussian process centered at the true stimulus frequency. Bottom: If a trial is judged to contain only signal tones that were drawn from the low- or high-frequency signal Gaussian distributions, the psychometric curve becomes sigmoid, as in a traditional 2AFC Task. (D) The Random Guess Model considers those participants that assign category identities at random, as if all tone bursts were drawn from the distractor distribution. Top: Given a particular category, the perceived probability of a certain tone frequency is flat across the stimulus range. Middle: The sensory process of the Bayesian observer is modeled as a Gaussian process centered at the true stimulus frequency. Bottom: If a trial is judged to contain only distractor tones, the psychometric curve becomes flat, as each tone is not informative to the category decision. Blue: p(v|C=L), red: p(v|C=H), purple: p(v|D).

However, in the presence of distractors, we expect participants’ performance to be substantially different. Because tones whose frequencies are far from the category-specific Gaussians are more likely to be distractors than signal tones (Figs 4C, D), participants may assign lower relevance to those tones. This effect would result in poorer accuracy for tones lying at the tails of the Gaussian distributions (Fig. 4D).

We developed a general framework to capture potential strategies that a participant could use when categorizing the three-tone sequences ^41,50–52^. These strategies depend on a participant’s internal model of stimulus relevance of the three tone frequencies and can be formalized using the Bayesian approach (Fig 5A). The full Bayesian model reflects a decision-making strategy for a participant (Fig 5A table, top). This model estimates the participant’s belief about the relevance of each of the three tones and integrates information across the tones accordingly. It also captures the participant’s prior beliefs of the trial categories. As alternative models, we considered two reduced variants of this model. In the no-distractor model, the participant assumed that all three tones were ‘signal’ (Fig 5A table, center). Conversely, in the random-guess model, the participant assumed that all three tones were irrelevant ‘distractors’ and made their choice randomly akin to a coin flip based on only their learned prior beliefs (Fig 5A table, bottom). This model controls for the possibility that participants ignored all the tones when they made their decision. Henceforth, we interpret our experimental findings using this three-model Bayesian framework.

**Fig. 5:**
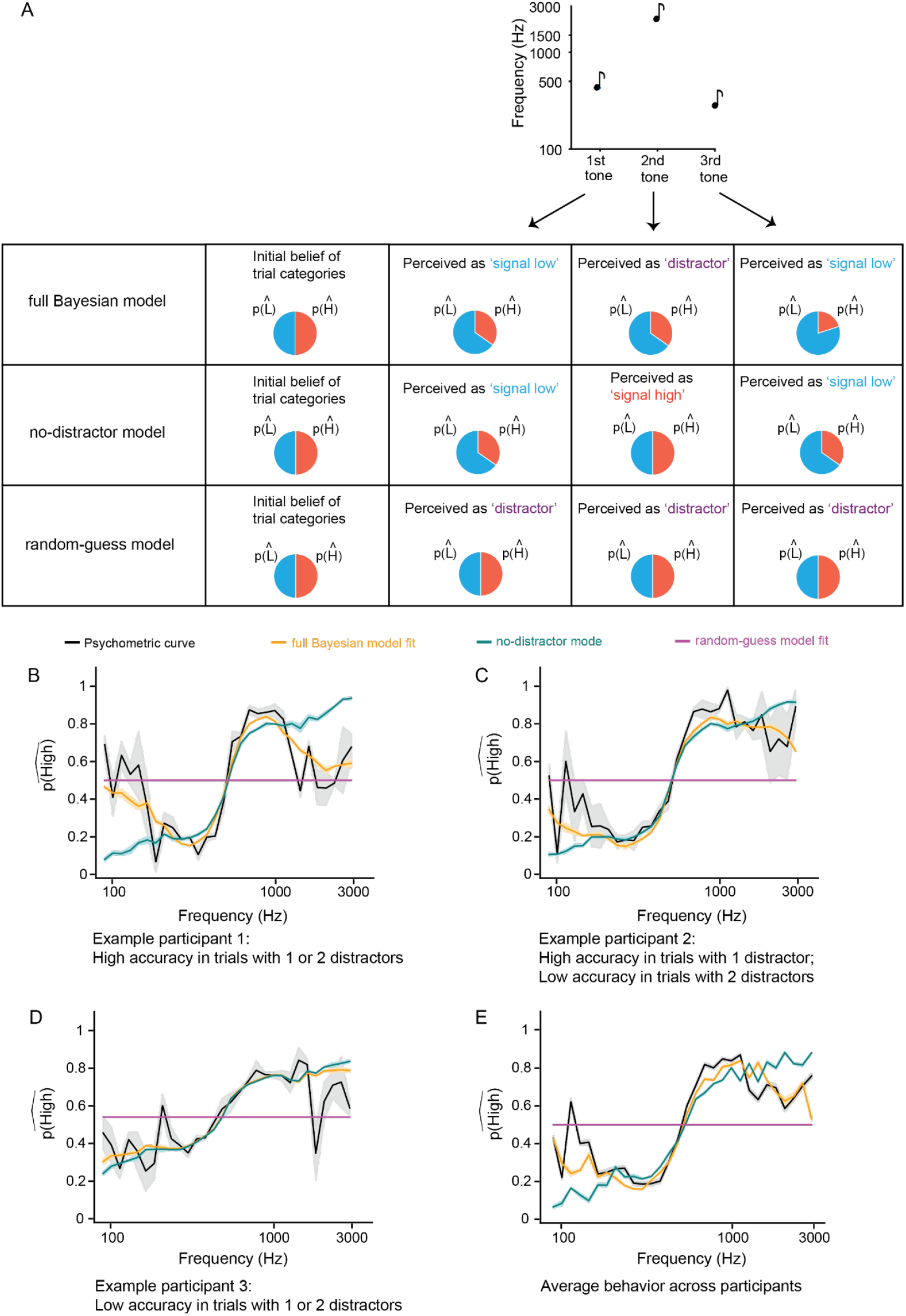
Schematic diagram of the 3 decision-making models; psychometric curves and model fits shown for both representative participants as well as averaged across all participants. (A) A schematic of all 3 decision-making models. (top) The full Bayesian model: the participant considers that any tone can probabilistically either be ‘signal’ or ‘distractor’ (see main text for more information). Using a hypothetical trial, we illustrate that deciding that a tone is ‘signal’ versus ‘distractor’ shifts participants’ category choice probabilities. (center) The no-distractor model: the participant considers all tones to be ‘signal’ tones, and thus a tone that the full Bayesian model interprets as a distractor might instead by interpreted as a high signal tone. (bottom) The random-guess model: the participant considers all tones to be ‘distractor’ tones and thus makes their decision randomly. 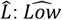-, 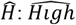 - category decision. (B, C, D) Psychometric curves (black lines) and respective model fits (full Bayesian model: orange lines, no-distractor model: teal lines, random-guess model: magenta lines) for 3 example participants (for posteriors of the full Bayesian model, and for GLM analysis, see Fig S3; for more participants see Fig S4). The x-axes of the psychometric curves span the frequency range of the stimulus tones, and the y-axes denote the mean probability of a 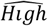 category choice response. Error bars: standard error of the mean. (B) Example participant with high task accuracy on trials with either one or two distractor tones. (C) Example participant with high task accuracy on trials with one distractor, and poor accuracy on trials with two distractors. (D) Example participant with poor accuracy on trials with both one distractor and two distractor trials. (E) Psychometric curve and model fits averaged across all participants.

We first examined the performance of a representative participant with high accuracy on trials with distractors (Fig 5B, one-distractor accuracy 80.59% and two-distractor accuracy 74.14%). Tones that were far from the two category distributions and thus more likely to be distractors were correlated less strongly with behavioral performance than tones closer to the middle of the distributions (Fig 5B, black). Additionally, the full Bayesian model (Fig 5B, orange) fit the participant’s nonmonotonic behavior considerably better than the no-distractor or random-guess models (Fig 5B, teal and magenta, respectively; Table 1 – two-sided Wilcoxon signed-rank test). These analyses collectively demonstrated that this participant used the relevance of the different tones when categorizing trials.

We next studied the performance of two participants with intermediate and low accuracy. Their choice probability for tones drawn from the flanks (highest or lowest frequencies) of the signal distributions changed less in comparison to the participant with higher accuracy, resulting in psychometric curves that were more sigmoidal in nature (Figs 5C, D). Furthermore, the difference in accuracy on trials with one and two distractors was lower for the low-accuracy than high-accuracy participants (Fig 5C: one-distractor accuracy 78.24% and two-distractor accuracy 59.48%, Fig 5D: one-distractor accuracy 59% and two-distractor accuracy 61.21%). Even for these participants, although the effect was smaller, the full Bayesian model provided a better fit to the psychometric curves compared to the other two Bayesian models (orange curve versus the teal and magenta curves; Table 1 – two-sided Wilcoxon signed-rank test).

These three examples suggest that potentially all the participants, irrespective of their accuracy, are impacted by stimulus relevance when choosing the stimulus category, albeit to varying degrees. To test this hypothesis across the entire population, we plotted the average psychometric curve and the average model fits (Fig 5E). We found that there the data were well fit by the full Bayesian model, which qualitatively illustrates a population-wide effect of stimulus relevance on category choice decisions.

Next, we quantified this effect by comparing the statistics for the fits of the Bayesian models. Because the number of fitting parameters differs across the three models (6 in the full Bayesian, 5 in the no-distractor, and 2 in the random-guess model (see Methods and Fig S5)), we used Bayesian Information Criterion (BIC) scores to compare them. A lower BIC score indicates a better model fit. For all participants, the full Bayesian model was better than the random-guess model (Fig 6A; population statistics: two-sided Wilcoxon signed-rank test on BIC, Z = -6.51, p = 7.55e-11) at describing categorization behavior. This implies that participants do not simply rely on their prior beliefs when classifying the tone sequences.

**Fig. 6:**
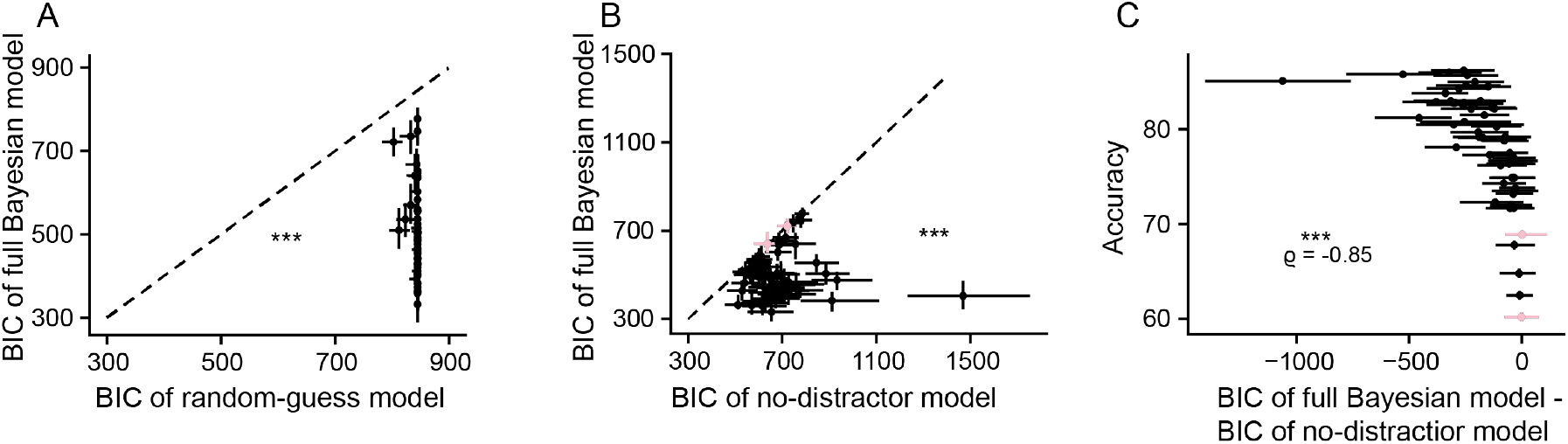
Nearly all participants were sensitive to stimulus relevance. On *unbiased* trials, BIC scores of the full Bayesian model were lower than that of the (A) random-guess model for all participants and (B) the no-distractor model for all except two participants. The two outliers are shown in light pink. Statistics: two-tailed Wilcoxon signed-rank test comparing model fits across the population. (C) Participants’ accuracy was inversely correlated with difference in BIC scores between the full Bayesian and no-distractor models. Statistics: Spearman correlation. In all panels, data point: one participant; ^†^*p*<0.05; \**p*<0.01, \*\**p*<0.001, \*\*\**p*<0.0001. Error bars in (A, B) and horizontal error bars in (C): 95% confidence intervals derived from respective model fits to 100 bootstrapped samples. Vertical error bars in (C): standard error of the mean, obtained by assuming that accuracy follows a binomial distribution; vertical error bars not large enough to be distinguishable.

For most participants, except for two, the full Bayesian model was also better than the no-distractor model (Fig 6B; same test, Z = -6.49, p = 8.41e-11). For the two outlier participants (Fig 6B, C; light pink), the full Bayesian and no-distractor model fits were similar (Table 1 – two-sided Wilcoxon signed-rank tests), indicating that they did not necessarily account for relevance when making decisions. However, for the rest, the full Bayesian model reproduced the main patterns in the data in an absolute sense because it accurately captured participants’ responses across tone frequencies (Figs 5B-E, Figs S4A, D, G). These results confirm that nearly all participants took stimulus relevance into account when making category decisions.

Finally, we correlated model fits to participants’ task accuracy. Across the population, the difference in BICs of the full Bayesian and the no-distractor models was inversely correlated with accuracy (Fig 6C, Spearman, ρ = -0.85, p = 8.53e-17). For highly accurate participants, the full Bayesian model performed particularly well. These data, once again, suggest that participants’ behavioral performance was determined by their ability to discriminate tones based on their decision relevance.

We also explored the space of non-Bayesian models that had fewer assumptions. For the first model, we implemented a simple boundary model that had three parameters: (1) the frequency dividing the low distractor range from the low-signal range, (2) the frequency dividing the low and high signal ranges, and (3) the frequency dividing the high signal range from the high-distractor range. We found that this model was informative for the high-accuracy participants and consistent with the full Bayesian model (Figs S2C, D), but was not informative for the other participants.

For the high-accuracy participants, this boundary model provided insight into the extent to which they discounted extreme stimuli. We found that these participants exaggerate the signal-frequency ranges by underestimating the lower signal boundary (separating distractor from low signal) and overestimating the upper signal boundary (separating high signal from distractor) (Fig S2A). This matches the full Bayesian fitting results, which also show a tendency to exaggerate the distance of the perceived generative signal distributions from the center of the stimulus range (Figs S5B, C).

For the second model, we implemented a generalized linear model (GLM) (Figs S3A, E). We found that this model was consistent with the full Bayesian model for high-accuracy participants. The observation that high- and intermediate-accuracy participants are impacted by stimulus relevance in their decision-making process was substantiated by the corresponding boundary models and GLMs (Figs S2D, E, S3F, G). The GLM analysis was also able to substantiate the finding that the difference in accuracy on trials with one and two distractors was lower for the low-accuracy than high-accuracy participants (Fig S4).

However, the boundary model and GLM differed in their ability to capture the behavior of low-accuracy participants. For low-accuracy participants, the boundary model assigned internal boundaries to these participants that caused most of the stimulus range to be labeled as distractor tones (Fig S2B). In contrast, the GLM results indicate that even these participants understood the generative distributions (Fig S3F), which is backed by the full Bayesian results showing that these participants were still best fit by the full Bayesian model. (Figs 5D, S3C). Although the GLM was able to describe the behavior of low-accuracy participants, it was not as informative as the Bayesian models regarding the interpretation of the parameters (Figs S3E-G). Together, these results indicate that individual variability in performance is not captured by the boundary and GLM models which primarily are driven by representations of the generative distributions. Therefore, other factors likely account for individual variability in performance.

### Sensory uncertainty and estimated task relevance underlie inter-participant variability

Individual participants’ ability to discriminate between generative distributions of the signal and distractor tones may depend not only on their subjective estimates of the tones’ task relevance (or more specifically, their estimate of the probability of distractors) but also on their measure of stimulus uncertainty (which is the standard deviation of the two Gaussian distributions) and their sensory uncertainty. For instance, some participants may have different prior convictions about the number of distractors (e.g., some may spend more time in noisy environments than others), which could bias how they integrate information over the three tones. Additionally, we expect that participants with low stimulus and sensory uncertainty would distinguish better between signal and distractor tone frequencies than participants with high uncertainty. A participant’s sensory uncertainty determines the variability in perception of the same tone frequency over multiple presentations, whereas the stimulus uncertainty captures the participant’s estimate of the variability in the underlying source generating the same stimulus. Thus, a participant with low stimulus and sensory uncertainties would perceive repeated presentations of the same tone more consistently and would estimate the distribution of the low (high) tone source to be narrower than a participant with high stimulus and sensory uncertainties. Consequently, we conjectured that estimated priors for the probability of distractors, stimulus uncertainty, and sensory uncertainty would drive individual participants’ category choice and task accuracy contributing to inter-individual variability.

To test the effect of these three factors on category choice, we first analyzed the inter-participant variability in these factors. We found that across the 56 participants, stimulus uncertainty (denoted by model parameter *σ*) was nearly constant (Fig S5D), whereas sensory uncertainty (denoted by parameter *σ*_*sensory*_) and the estimation for the probability of distractors (denoted by model parameter *p*_*distractor*_) were participant-specific (Figs S5A,E,F). To quantify a relationship between *p*_*distractor*_ and *σ*_*sensory*_, we defined a metric ‘sigmoidicity’ (Fig 7A; see Methods) which quantitatively captured participant’s category decisions across all trials. We found that this metric was significantly influenced by *p*_*distractor*_ and *σ*_*sensory*_ as well as by the interaction between these two parameters (Fig 7B; Table 1 – two-factor regression model). Specifically, sigmoidicity increased with *σ*_*sensory*_ and decreased with *p*_*distractor*_. In other words, sigmoidicity could be used to quantify the extent to which the participants considered the presence of distractors when categorizing.

**Fig. 7:**
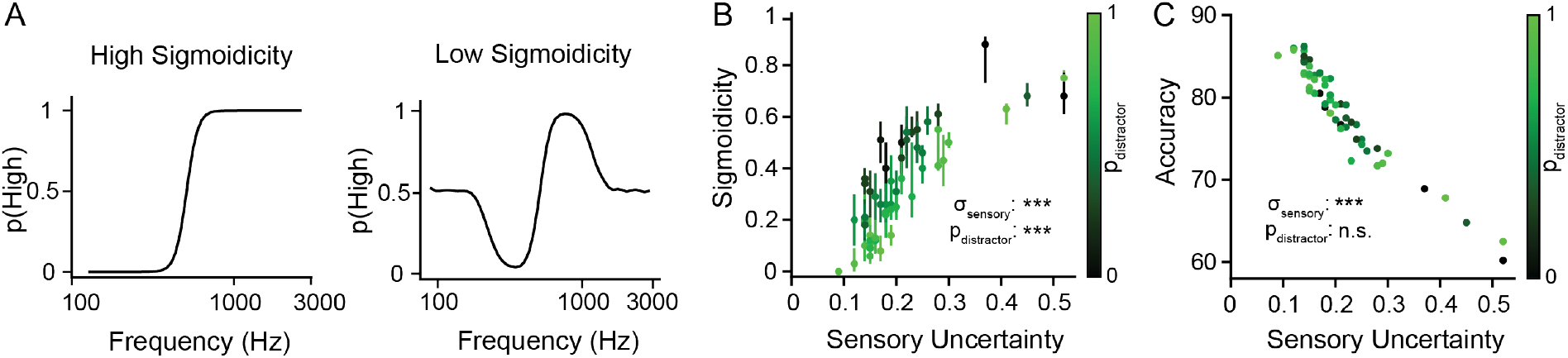
For an individual participant their category choice was driven by both their sensory uncertainty and their subjective estimates of the probability of distractors, but their accuracy was mostly driven by their sensory uncertainty. (A) Cartoon models of different psychometric curves illustrate the qualitative range of sigmoidicity. Left: high sigmoidicity, right: low sigmoidicity. (B) Each data point corresponds to a single participant; color denotes *p*_*distractor*_ which is the participant’s estimated probability of distractors (see color bar); Black: *p*_*distractor*_ ∼ 0; Green: *p*_*distractor*_ ∼ 1. *p*_*distractor*_ is a fitting parameter in the full Bayesian model and has value 0 in the no-distractor model. The model parameter *σ*_*sensory*_ captures participant’s sensory uncertainty. Statistics: effect of *p*_*distractor*_ and *σ*_*sensory*_ on sigmoidicity using a two-factor regression model. (C) Participants’ accuracy did not significantly depend on *p*_*distractor*_ but was inversely correlated with their sensory uncertainty. *p*_*distractor*_ is color coded as in (B). Statistics: effect of *p*_*distractor*_ and *σ*_*sensory*_ on accuracy using a two-factor regression model. In all panels, data point: one participant; ^†^p<0.05; *p<0.01, **p<0.001, ***p<0.0001. Error bars in (B) and horizontal error bars in (C): 95% confidence intervals derived from respective model fits to 100 bootstrapped samples. Vertical error bars in (C): standard error of the mean, obtained by assuming that accuracy follows a binomial distribution; vertical error bars aren’t large enough to be distinguishable.

We then tested the effect of *p*_*distractor*_ and *σ*_*sensory*_, on participants’ accuracy. Although we did not find a relationship between accuracy and *p*_*distractor*_ (Fig 7C; Table 1 – two-factor regression model), we found a strong, roughly linear, relationship between accuracy and *σ*_*sensory*_ (Spearman, ρ = -0.96, p = 3.15e-31). Taken together, these data demonstrate that participants had a similar measure of stimulus uncertainty but differed in their sensory uncertainty and estimate of relevance. Further, although participants’ category decisions were shaped by both their estimate for the probability of distractors and their sensory uncertainty, much of their differences in accuracy may be driven by differences in their ability to distinguish between the tone frequencies.

Next, we tested whether listeners differentially weighed tones based on their position in the sequence when making their category decision. To test this idea, we developed an alternative Bayesian model with a nonuniform psychophysical kernel in which we substituted *p*_*distractor*_ for 3 tone-position dependent versions of *p*_*distractor*_. We found that participants weighed the first two tone positions 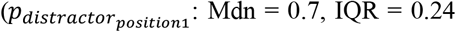; 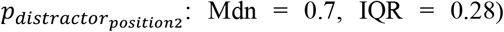 equally but slightly more than the third tone position 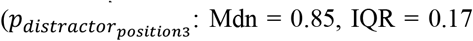 Fig S6G; Table 1 – one-way rmANOVA, post hoc two-sided Wilcoxon signed-rank test) when making their category decision. Additionally, values for 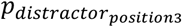 but not 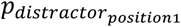 and 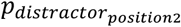 were statistically different than *p*_*distractor*_ (Fig S6G; Mdn = 0.59, IQR = 0.43; Table 1 – two-sided Wilcoxon signed-rank test). The influence of all three position-dependent weights on sigmoidicity and accuracy was consistent with results (Figs. S6A-F; Table 1 – Spearman) from the full Bayesian model (Figs. 7B, C), in that both *p*_*distractor*_ and *σ*_*sensory*_ were correlated with sigmoidicity, while *σ*_*sensory*_, but not *p*_*distractor*_, was correlated with accuracy. The finding that the third tone position is discounted relative to the first two tones is also supported by results from the simpler GLM model (Fig. S6H; Table 1 – one-way rmANOVA, post hoc two-sided Wilcoxon signed-rank test). We found that this alternative tone location-dependent Bayesian model outperformed the location-independent model, likely because it was able to capture this discounting of the tone in the third position (two-sided Wilcoxon signed-rank test on BIC, T = 0, p = 7.55e-11). Overall, these data show that participants may have used information from all three tone positions but weighed the third tone somewhat less than the first two.

Using this differential weighting of the tone positions, we reconstructed the psychometric curves (Fig 8) for the representative participants shown in Fig 5. This reconstruction accounts for a participant’s accumulation of evidence across the three tone positions. Interestingly, we found that although the full Bayesian model was simpler, both models adequately fit the experimental psychometric curves and for both the models the underlying p_distractor values were higher than the true experimental value of 0.3. Participants thus had a tendency to ignore tones more than the statistics of the task would require.

**Fig. 8:**
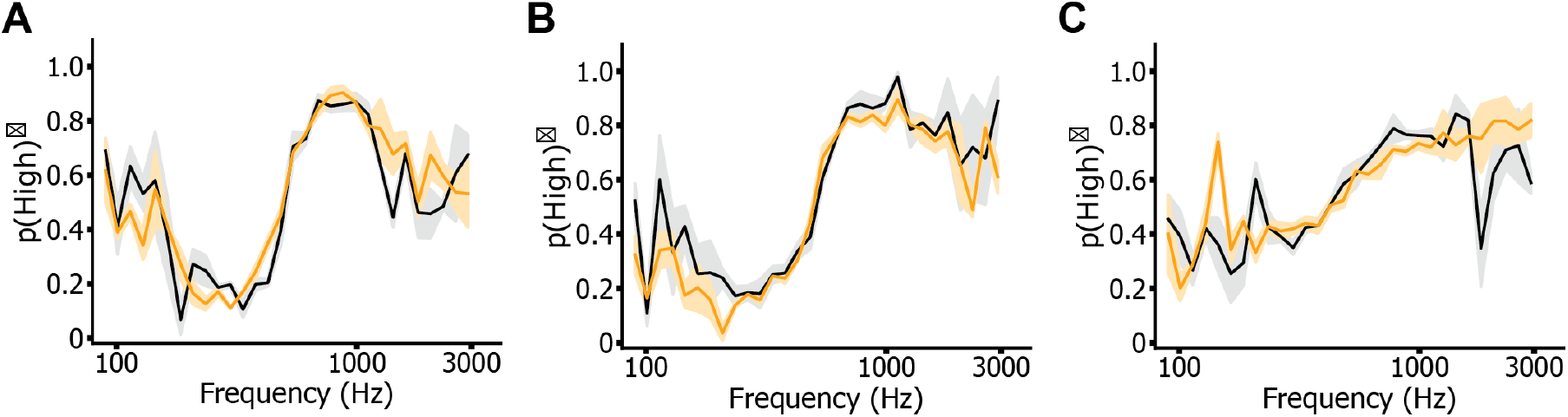
Psychometric curves for representative participants fitted using the tone-position dependent Bayesian model. (A, B, C) Psychometric curves (black lines) and model fits of the tone-position dependent Bayesian model (orange lines) for the same 3 example participants as Fig 5. The x-axes of the psychometric curves span the frequency range of the stimulus tones, and the y-axes denote the mean probability of a 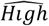 category choice response. Error bars: standard error of the mean. (A) Example participant with high task accuracy on trials with either one or two distractor tones. (B) Example participant with high task accuracy on trials with one distractor, and poor accuracy on trials with two distractors. (C) Example participant with poor accuracy on trials with both one or two distractors.

### Participants rely on past statistics of category probabilities

Having established how stimulus uncertainty, sensory uncertainty and stimulus relevance shape auditory categorization, we next asked whether and how participants’ learning of past statistics of the category probabilities informed their current decision. We first analyzed the influence of short-term statistics by computing for *unbiased* sessions the mean probability of a 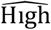 category choice response for a given trial (*t*) conditioned on the category of the previous trial (*t* − 1), 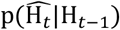 and 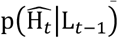 (Fig 9A). We found that the difference between the two curves was quite variable across participants, suggesting that although some participants’ decisions were substantially affected by the short-term history of category probabilities (Figs 9C, D), others’ decisions were not (Fig 9B). Overall, across the population, the category of the previous trial (i.e., short-term stimulus statistics) significantly influenced participants’ choices on the current trial (Fig 9E; Δ_*short*−*term*_: difference between 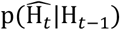 and 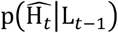 at central frequency of 511Hz; central frequency: frequency which is the mid-point of the experimental frequency range in log-space; two-sided Wilcoxon signed-rank test on Δ_*short*−*term*_, Z = 6.04, p = 1.58e-9). Moreover, we found that this influence was only restricted to the category of the previous trial and did not substantially depend on whether the participant responded accurately to the trial (Fig S7; Table 1 – two-sided Wilcoxon signed-rank test on Δ_CorrectVsIncorrect_). Combined, these data confirm the presence of category-driven short-term learning in decision-making.

**Fig. 9:**
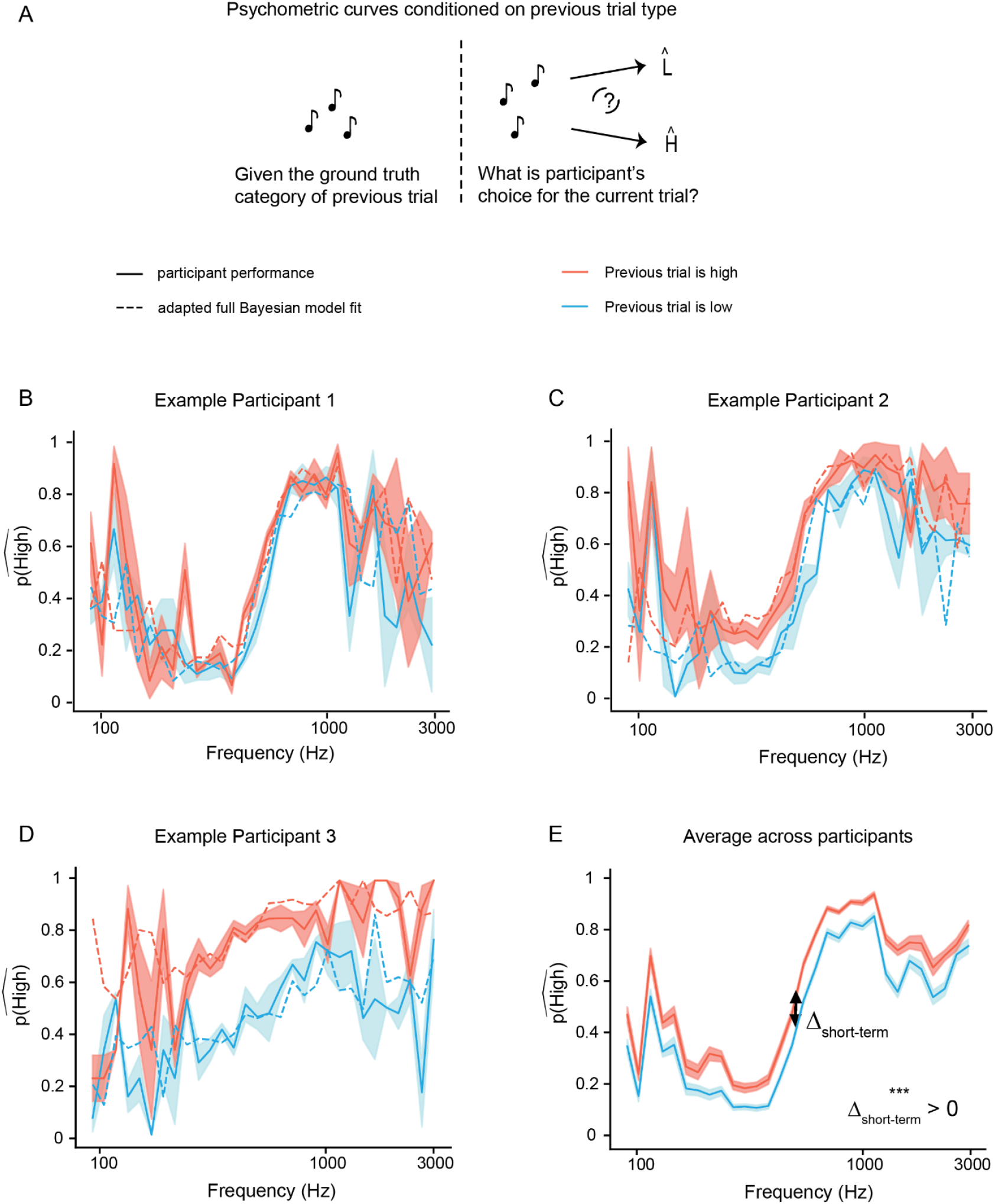
To a varying degree, participants incorporated short-term statistics of category probabilities in their decisions. (A) Schematic shows the effect of learning short-term statistics. (B-D) Average psychometric curves (solid lines-light blue: previous trial is low; red: previous trial is high) and the adapted full Bayesian model fits (same colors, dashed lines) conditioned on the previous trial’s category type for the same three participants as Figure 5 in the *unbiased* session. Error bars: standard error of the mean. Difference between 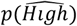 computed at the central tone frequency 511 Hz Δ_*short*−*term*_: (B) ∼0.1, (C) ∼0.22 and (D) ∼0.32. (E) Corresponding psychometric curves averaged across all participants in the *unbiased session*; Δ_*short*−*term*_: ∼0.12. Statistics: two-tailed Wilcoxon signed-rank test on Δ_*short*−*term*_. In panel (E), ^†^*p*<0.05; \**p*<0.01, \*\**p*<0.001, \*\*\**p*<0.0001.

In conjunction with short-term learning, participants may also use long-term learning when there is a bias in the trial categories toward low or high choices over a longer time scale. To test whether participants were affected by short- and/or long-term learning, in separate behavioral sessions, we changed the probability of the low-category trials. Specifically, *p*_*L*_ = 0.7 in *biased low* sessions and *p*_*L*_ = 0.3 in *biased high* sessions (Fig 11A; Participants: N = 53 in *biased low* and N = 48 jointly in *biased low* and *biased high*). Anecdotally, individual participants with high values of *p*_*distractor*_ did not use either type of learning when making decisions (Fig 10B; Fig S8A). However, participants with lower values of *p*_*distractor*_ seemed to use previous trial statistics to varying degrees (Figs 10C, D). Some participants used both short- and long-term learning (Fig 10C; Fig S8B), whereas others primarily used long-term learning (Fig 10D; Fig S8C). Overall, in the population, there was a clear effect of both the session type and the last trial on category choice (Fig 10E, Figs S7-9; Δ_*long*−*term*_: difference between psychometric curves from *biased low* and *biased high* sessions at central frequency; population statistics: two-sided Wilcoxon signed-rank test on Δ_*long*−*term*_, Z = 8.63, p = 6.34e-18; two-sided Wilcoxon signed-rank test on Δ_*short*−*term*_, *biased low*: Z = 5.94, p = 2.91e-9; *biased high*: Z = 5.94, p = 2.91e-9). Thus, data across the three sessions – *unbiased, biased low* and *biased high* – suggests that both, short- and long-term learning influenced participants’ decision-making, presumably depending on their relevance estimates.

**Fig. 10:**
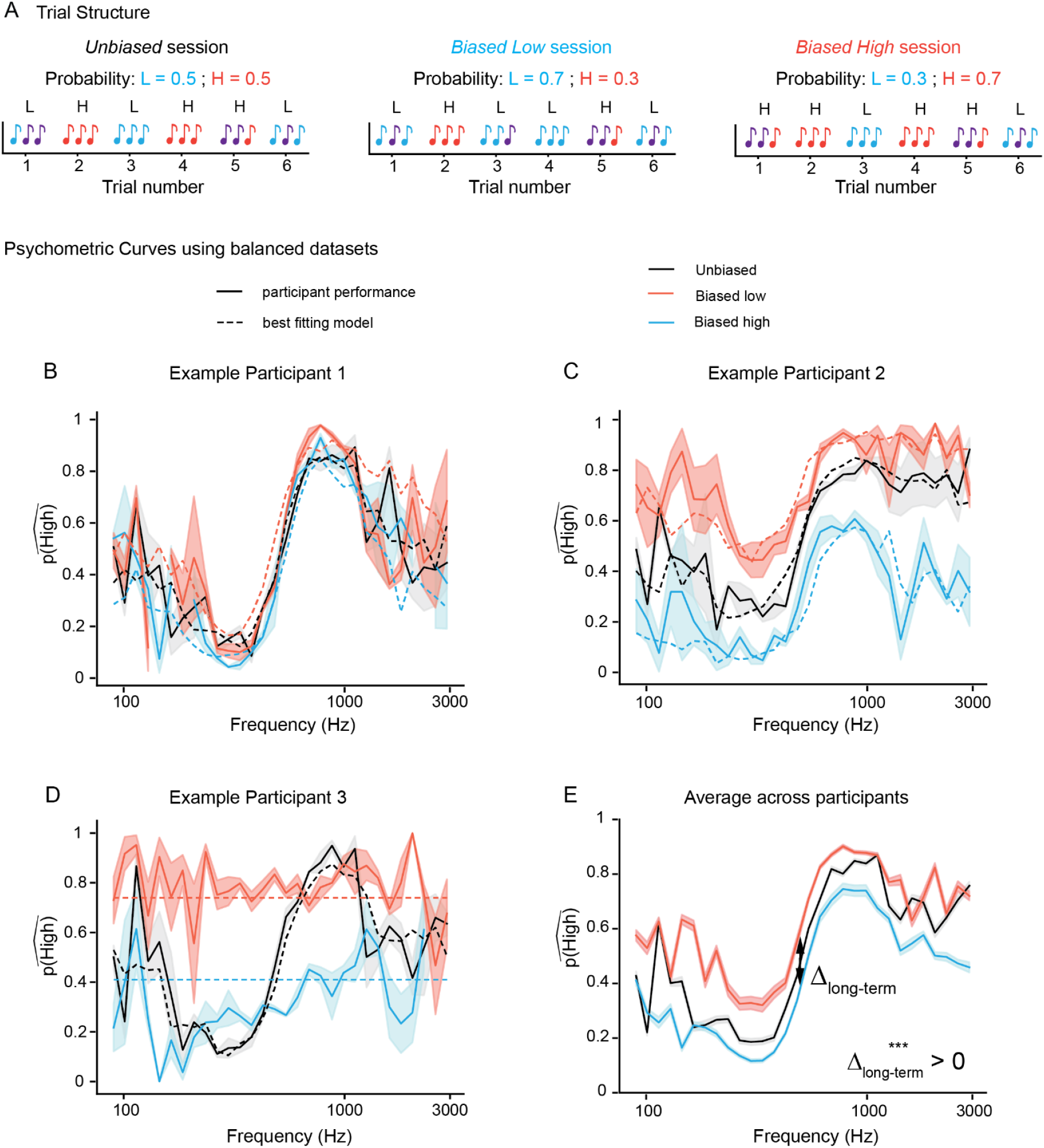
To a varying degree, participants’ decisions were affected by long-term statistics of category probabilities. (A) Example stimuli from *unbiased, biased low*, and *biased high* trials. Individual tones are color-coded according to their underlying distribution as shown in Fig. 1B. (B-D) Average psychometric curves for three participants in the *unbiased* (black), *biased low* (light blue) and *biased high* (red) sessions. Respective model fits also shown using dashed lines; (B, C) fit using the full Bayesian model and (D) fit using the random-guess model for the biased sessions and the full Bayesian model for the unbiased session. Error bars: standard error of the mean. Curves for the *biased* sessions are evaluated using balanced datasets (see Methods). Curves for the *unbiased* trials are similar to black lines in Figs 5B-D. Difference between 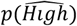 for *biased low* and *biased high* trials computed at the central tone frequency 511 Hz. Δ_*long*−*term*_: (B) ∼ 0.06, (C) ∼ 0.35 and (D) ∼ 0.51. (E) Psychometric curves averaged across all participants; Δ_*long*−*term*_: ∼0.2. Statistics: two-tailed Wilcoxon signed-rank test on Δ_*long*−*term*_. In panel (E), ^†^*p*<0.05; \**p*<0.01, \*\**p*<0.001, \*\*\**p*<0.0001.

**Fig. 11:**
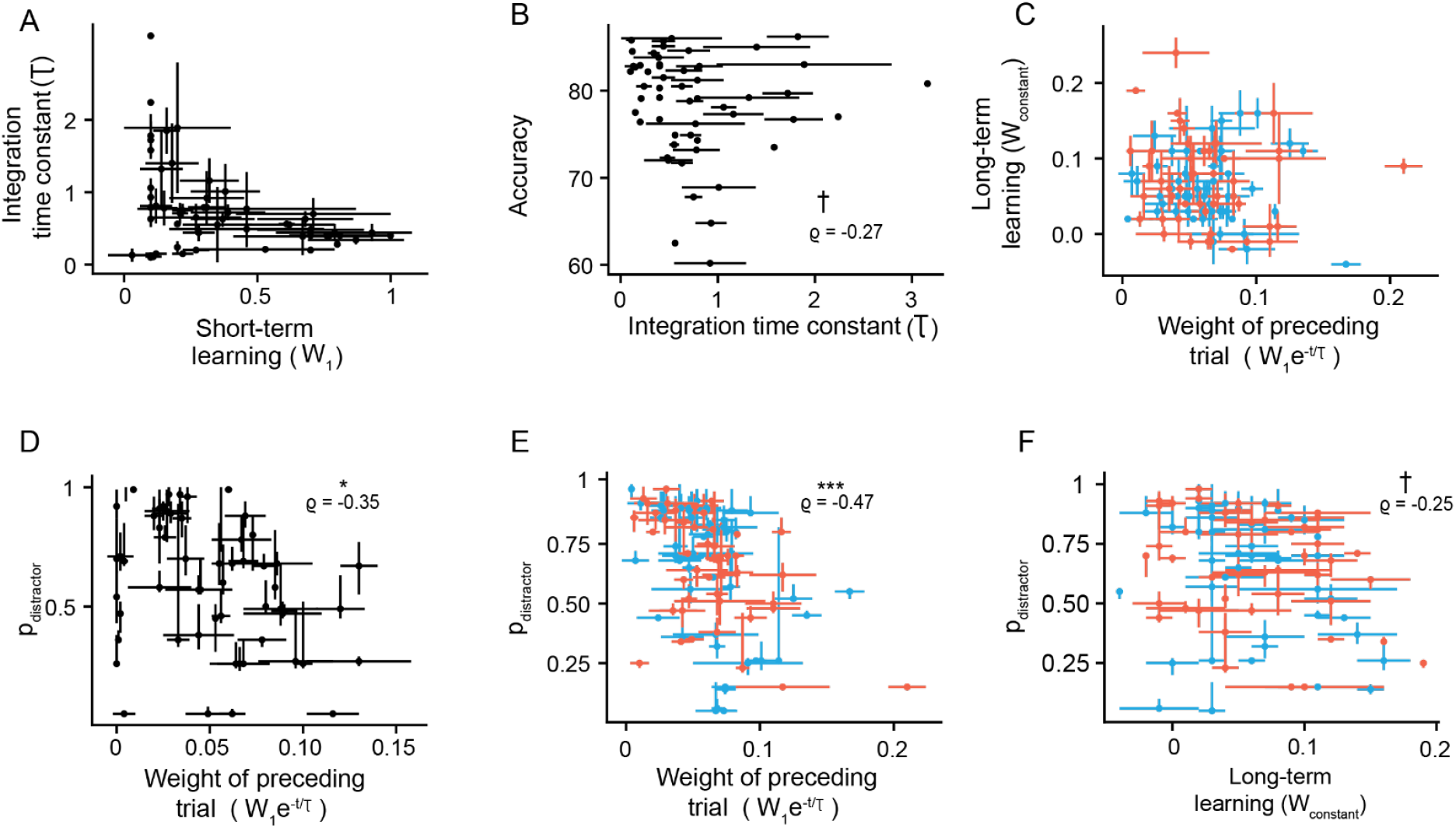
Participants subjective estimate of distractor probability varies inversely with their learning of stimulus statistics when making category decisions. (A) On *unbiased* trials, nearly all participants (black, N=55) exhibited short-term learning (*W*_1_ > 0). (B) Their accuracy was inversely correlated with the time constant (τ) for such learning. Further, the median number of previous trials over which participants’ retained information was 0.63, indicating that participants retained information from the immediately preceding trial. (C) On *biased* trials (*biased low*: light blue, N=53; *biased high*: red, N=48), the effect of short- and long-term learning varied substantially across participants. Across all three sessions, *p*_*distractor*_ was inversely correlated with (D, E) short-term learning given by *W*_1_*e*^−1/τ^ and in the *biased* sessions, it was also inversely related to (F) long-term learning captured using *W*_*constant*_. All panels, data point: one participant; ^†^*p*<0.05; \**p*<0.01, \*\**p*<0.001, \*\*\**p*<0.0001; Statistics: Spearman correlation; error bars: standard error of the mean.

To further quantify how learning modulates category decision-making, we adapted the full Bayesian model to incorporate both short- (exponential weighted: *W*_1_*e*^−*d*/τ^) and long-term (constant: *W*_*constant*_) components (Values of *W*_1_ and *W*_*constant*_ ∼ 0 implies that there was not any learning, whereas values ∼1 implies learning; τ is a time constant that captures the number of preceding trials that a participant may have used to learn the short-term stimulus statistics). This adapted model, thus, combined the effects of sensory uncertainty, stimulus relevance, stimulus uncertainty, as well as learning of long- and short-term stimulus statistics.

Using this framework, we asked how participants measure of stimulus relevance and their sensory uncertainty modulated their learning of stimulus statistics. Thus, we first tested whether participants accounted for stimulus relevance when categorizing trials during *biased low* and *high* sessions. We found that the performance of most participants, except for two, was best fit by the full Bayesian model (Fig S10). These two outlier participants primarily used their learned prior and their performance was thus best fit by the random-guess model (Figs S10A, B, D, E). These data reinforce our previous finding that stimulus relevance is central to categorization.

Next, we assessed the degree to which participants learned the short-term statistics of trials by fitting the adapted version of the full Bayesian model to data from the *unbiased* session. We found that nearly all participants demonstrated significant short-term learning (Fig 11A; *W*_1_ > 0 for N = 55). However, there was substantial variability across participants such that for some participants, *W*_1_∼0, whereas for other participants, it was ∼1. Similar heterogeneity was also observed in the time constant for learning such that the participants’ time constant was inversely correlated with their accuracy (Fig 11B; Spearman, ρ = -0.27, *p* = 0.042). However, across the population, the average time constant (τ) in the *unbiased* session was ∼1 trial (Mdn: 0.63 trial, IQR: 0.44; Figs 11A, B). The nature of *W*_1_ (Fig 11C) and τ (Fig S11; Table 1 – Spearman correlation with accuracy) was similar in the *biased* sessions. Jointly, these results indicate that although most participants show signs of short-term learning when making category decisions, this learning was largely restricted to the immediately preceding trial.

Similar to short-term learning, we also expected some participants to exhibit effects of long-term learning. Indeed, most participants learned the long-term trial statistics (Fig 11C), but this learning was not correlated to their short-term learning (Spearman; Table 1). Thus, participants appear to use both the bias in the session and their knowledge of previous trials albeit to different degrees when making decisions.

Next, we probed how these two learning mechanisms relate to participants’ estimated probability of distractors. We found that the weight of the preceding trial (*W*_1_*e*^−1/τ^) was inversely correlated with *p*_*distractor*_ for all three sessions (Figs 11D, E; Spearman, *unbiased*: ρ = -0.36, *p* = 0.0069, *biased low and high*: ρ = -0.47, *p* = 1e-6). Similarly, long-term bias (*W*_*constant*_) was also inversely correlated with *p*_*distractor*_ (Fig 11F; Spearman, *biased low and high*: ρ = -0.25, *p* = 0.013). This indicates that participants’ measure of relevance uncertainty was inversely correlated with their stimulus-category knowledge derived over both short- and long-time scales. This suggests that participants who are poor at differentiating between signal versus distractor tones likely rely on other sources of information such as past knowledge when making category choices.

Finally, we asked how the various parameters that drive performance – sensory uncertainty, stimulus relevance, stimulus uncertainty, and learning relate to task accuracy. We found that sensory uncertainty was a strong predictor of accuracy, accounting for much of the variance (Figs 7C, S12; Spearman, *biased low and high*: ρ = -0.75, *p* = 2.38e-19). Thus, the main factor in deciding how proficiently people categorized sounds could be interpreted as how well they perceived the different tone frequencies. These results support the basic assumption of Bayesian models that sensory uncertainty is a fundamental factor in decision making.

## Discussion

It has long been known that humans and other animals can isolate signals of interest from background in noisy, crowded environments. When presented with two competing auditory streams, participants’ neural responses to the task-relevant stimuli are enhanced^53,54^, but, their processing of the irrelevant distractor stimuli is mediated by attention^55,56^. Visual spatial and feature attention affects the neural representation of task-irrelevant sensory information^57,58^. Furthermore, because such information is often incompletely suppressed, the presence of distractors impairs human decision-making behavior^15^. However, much of this work has studied the effects of relevance in isolation. When making decisions in noisy, uncertain environments, humans often incorporate multiple sources of information. One such source is the long-term sensory statistics of the stimuli of interest^30,59–61^ – studies have found that sensitivity to these statistics introduces observable biases in perception. However, in addition to tuning to long-term regularities, our sensory representations often need to rapidly adapt to short-term changes^62^. Indeed, such adaptation underscores context-dependent speech perception and categorization^27^. In our work, we investigated this complexity in perceptual decision-making by testing how participants’ sensory uncertainty, measure of stimulus relevance, stimulus uncertainty, and learning interacted to shape their auditory categorization behavior.

We found that participants differentially combined these factors when making category decisions. Specifically, when categorizing tone sequences consisting of both category-irrelevant (distractors) and relevant tones, participants weighed individual tones by their task relevance (Figs 2,3; Figs S1-4,10). Additionally, nearly all participants had similar stimulus uncertainty but diverged in their sensory uncertainty and their estimates of stimulus relevance (Fig S5). In fact, participants’ psychometric curves and category choices were jointly determined by both their sensory uncertainty and their subjective estimate of stimulus relevance, whereas their accuracy was mostly driven by their sensory uncertainty (Fig 7). We also observed that participants’ prior expectations of stimulus category were affected by their knowledge of stimulus statistics over both short and long timescales and this effect was inversely correlated with participants’ estimated probability of distractors (Figs 9-11; Fig S7-10). Additionally, we treat stimulus uncertainty as an estimate that the participants make about the stimulus distribution. However, it is possible that some participants do not explicitly separate the estimates of the underlying stimulus distributions and internal noise.

### Flexibility of experimental paradigm

Compared to previous work^30,63^, our experimental paradigm is unique in that we could modulate the multiple factors of sensory uncertainty, stimulus relevance, stimulus uncertainty and learning all within the same stimulus sequence. For example, because our stimulus was a three-tone sequence and because we could manipulate the number of task-irrelevant distractors (Fig 1), we could examine not only how participants accounted for relevance during categorization but also how their sensory uncertainty might have affected their associated measures of relevance (Figs 2,5-7; Fig S1). Additionally, although humans use both short- and long-term knowledge to efficiently make decisions^26,33^, most studies probing the influence of learned expectations on categorization have largely examined either short- or long-term effects without testing for their interaction^64–66^. Because in part of our experiment we biased the tone sequences towards either the low or the high category, we could investigate how participants’ reliance on long-term stimulus statistics interacted with their learning of short-term statistics i.e., the effect of the previous trial’s category (Fig 11). Thus, our experimental setup allowed us to simultaneously probe multiple factors that often underlie category decision-making. However, a potential shortcoming of our experimental design is that we did not vary stimulus uncertainty. Future studies could examine how this parameter interacts with all the other behavioral factors when making category decisions. In addition, we treat stimulus uncertainty as an estimate that the participants make about the stimulus distribution. However, it is possible that some participants do not explicitly separate the estimates of the underlying stimulus distributions and internal noise.

### Bayesian approach to study variability and interplay of relevance and uncertainty

To quantify the interplay between all these factors, we leveraged a Bayesian model (Figs 1,5-11). Bayesian models are a powerful tool that has been successfully used to study human behavior in sensorimotor learning^12,67^, sensory perception^25,68^, categorization^30,42^ etc. Our Bayesian analysis allowed us to characterize how individual participants may vary in their categorization behaviors. Across all three sessions, performance of nearly all participants was best captured by the full Bayesian model (Figs 6,11; Figs S4,10). However, in the *unbiased* session, few participants were found to be equally well fit by both the full Bayesian and the no-distractor models, suggesting that they did not take relevance into account when making category decisions. In the *biased* sessions, two participants who relied only on long-term stimulus statistics to make their decisions were best captured by the random-guesses model (Fig S10B, E).

In addition to this heterogeneity across strategies, we found that even among the participants best-captured by the full Bayesian model, there was substantial variability (Figs 7,11; Figs S2-6,8) resulting from participants’ sensory uncertainty, their subjective measure of relevance, and their degree of short- and long-term learning. For instance, by modeling sensory uncertainty using the parameter *σ*_*sensory*_, we found that although, on average, the value of participants’ sensory uncertainty is consistent with previous literature^30^, some participants have low sensory uncertainty and high task accuracy whereas some others have substantially high sensory uncertainty and poor task accuracy (Fig 7). This variability also underscored participants’ measure of stimulus relevance and their learning of stimulus statistics. In the latter, we observed that not all individuals used their prior information. Overall, these results illustrate diversity in the human perceptual decision-making process and follow prior studies that have elaborated upon individual differences in performance across several speech and non-speech tasks ^69,70^. Additionally, because our fitting procedures for the Bayesian models suggest that the participants are likely veridical about the stimulus features, their variability in categorization behavior mostly results from the cumulative effects of internal processes such as determination of stimulus relevance, sensory uncertainty and reliance on short- and long-term learning. This finding is consistent with previous work that attributes individual differences in the cocktail-party effect to stimulus-independent, internal stochastic processes ^71,72^.

Future work could use similar Bayesian approaches to probe whether in such auditory categorization tasks, participants use the same strategy throughout the experiment or switch between different strategies^73^ based on the complexity of the trial. Further, we conjecture that given the interdependence of stimulus relevance and strategy, it is likely that our analysis is a simplification of human behavior, and that relevance may be a time-varying property of participants. This contrasts with sensory uncertainty, which is likely to be a static parameter intrinsic to a given participant.

### Learning

The Bayesian analysis also allowed us to explore how the different forms of uncertainty – sensory uncertainty, stimulus relevance and stimulus uncertainty – interact with learning of stimulus statistics during categorization. We found that both short-term and long-term learning biased participants’ behavior in a manner consistent with previous studies ^30,74^. Moreover, these effects were independent (Fig 11) of each other^60^, but they were both inversely correlated with the relevance parameter *p*_*distractor*_. In other words, when making category decisions, participants combined their uncertainty about stimulus relevance with other sources of information^75^. Additionally, because the influence of long-term learning in the experiment grew with time (Fig S9), participants may have adapted to the underlying stimulus statistics. Future work should investigate the neural mechanisms that drive the dynamics of this adaptation, over both short and long timescales, and how such adaptation may be altered by the presence of distractors.

### Future directions

The interplay between multiple factors might also have implications for the broader understanding of how humans navigate real-life crowded noisy environments. For example, spatial cues^76,77^ and attention^78–80^ are integral to parsing multiple sound sources, but it is not yet well understood how humans can effortlessly identify relevant sounds in these noisy conditions. In our study, participants categorized three-tone sequences into high or low categories such that their category decisions were not greatly affected by tone positions (Fig S6). However, real-life stimuli such as speech^81^ and musical chords have rich information both within and across sequences. Whether such structure can aid relevance discrimination remains untested. Furthermore, our results indicate that participants’ estimates of stimulus relevance varied inversely with their learning of stimulus statistics over different time scales. Future work should explore the role of relevance in real-life scenarios and whether prior knowledge of one’s environment may detract from identification of relevant sounds. By covering many aspects of the complex process of auditory category learning, our approach may also open new ways of studying the neural basis of audition. Short- and long-term learning^82–86^, uncertainty in relevance of stimuli^57^, uncertainty in category of trials^87^ and sensory uncertainty may all be captured by distinct circuit elements and brain regions^88,89^. As such, experiments that vary these factors in a model-driven framework can enable new approaches towards understanding neural computation in the auditory system.

## Methods

### Ethics Statement

The experimental protocol was approved by the Institutional Review Board of the University of Pennsylvania. All procedures were carried out in accordance with the IRB guidelines. All participants participated voluntarily and provided informed consent in writing prior to participating.

### Human Psychophysics Apparatus

We used the crowdsourcing online platform provided by Prolific to recruit participants. We collected participants’ consent and demographic data using Qualtrics and conducted all experiments using Pavlovia. All subjects reported having normal or corrected-to-normal hearing. Subjects were required to use a laptop or desktop computer to complete the experiment. Each participant had to pass a ‘headphone-check’ experiment that helped ensure that they were wearing headphones or earphones as instructed^90^ before continuing with the experiment. This headphone-check required participants to judge which of three pure tones is the quietest and is designed to screen out non-headphone users due to phase-cancellation.

### Stimulus creation and task design

Human participants performed an auditory categorization task in which they reported the frequency category (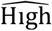 or 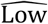) of a three-tone sequence (each tone is a sinusoid; tone duration: 300 ms during training, 280 ms during testing; inter-tone interval: 175 ms during training, 165 ms during testing; 44100 Hz sampling rate; 20 ms on- and off-ramps). Longer duration and intervals were used during training to give the listener more information about the task. Once they had learned the task structure, these parameters were reduced to reduce overall experiment duration. Each of the three tones *v* probabilistically could be either signal or distractor; that is a signal tone was relevant to the category membership of a sequence, whereas a distractor tone was irrelevant. The signal frequency probability distributions were Gaussian (*p*_*G*_; Eqs. 1a, b) in log(Hz) space: the mean of the low-frequency distribution (*µ*_*L*_) was 2.55, the mean of the high-frequency distribution (*µ*_*H*_) was 2.85, and standard deviation of both (*σ*_*expt*_) was 0.1. In Hz, the mean frequencies for the low and high distributions correspond to 355 Hz and 708 Hz respectively. The distractor frequency probability distribution was uniform (*p*_*U*_; Eq. 1c) and overlapped with both Gaussian distributions. For the distractor probability, the frequencies ranged from a = 90 Hz to b = 3000 Hz.

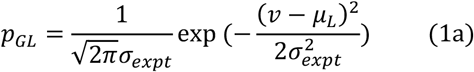

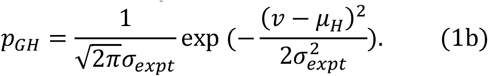

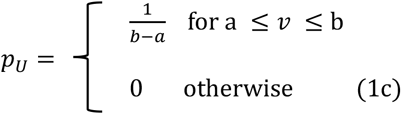

Overall, a low (high) category sequence was a combination of low-frequency (high-frequency) signal tones and distractor tones. The complete frequency range of the tones was between 90 – 3000 Hz and was uniformly sampled in log 10 space such that the frequency of each tone was one of 30 possible values. Because of the discrete nature of the frequencies, the equations presented here are probabilities across the 30 possible frequency values rather than continuous probability density functions.

Participants completed three sessions of the experiment: *unbiased, biased low*, and *biased high*. Tone sequences in the *unbiased* sessions were equally likely to be drawn from either the high or low categories, i.e., *p*_*L*_ = 0.5. Conversely, in the *biased low* (*biased high*) sessions, we overrepresented the corresponding distribution such that *p*_*L*_(*p*_*H*_) = 0.7. Across all three sessions, the probability that each tone was signal was *p*_*S*_ = 0.7 and of it being a distractor was *p*_*D*_ = 0.3. This meant that, of the set of three tones, the probability that the set included just one distractor was 3(0.3)(1 - 0.3)(1 - 0.3) = 0.44. The signal and distractor tone probabilities were chosen based on pilot data internally collected. Specifically, for *p*_*D*_, we tested values of 0.2, 0.3 and 0.4 and chose *p*_*D*_ = 0.3 based on a tradeoff between participants’ accuracy and their self-reported difficulty of the experimental task.

The generative process of the experimental stimuli can be summarized as,

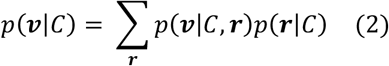

where ***v*** is the set of three experimental tones frequencies *v*_1_, *v*_2_, *v*_3_ (in units of log(Hz)). ***r*** is the set of tone relevance indicators *r*_1_, *r*_2_, *r*_3_ ∈ {*R, I*} such that *r*_*d*_ = R implies that the tone *v*_*d*_ is signal (relevant) and *r*_*d*_ = I implies that it is a distractor (irrelevant). *C* is the category of the sequence (high (*H*) or low (*L*)). Because the tones and their relevance are conditionally independent, given a trial’s category, we can rewrite Eq. 2 as:

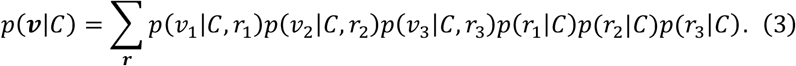

For example, in a high-category sequence, for a given tone *v*_*d*_,

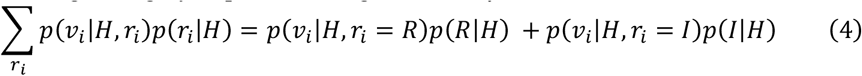

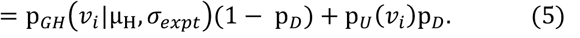

The priors in the generative process are *p*(*C* = *L*) = *p*_*L*_ and *p*(*C* = *H*) = *p*_*H*_ = 1 − *p*_*L*_.

### Participants

In total, 48 healthy adults (22 female; mean ± SD age of the 48 adults: 26.32 ± 7.17) successfully completed the full study. Of the 169 adults who signed up for the study, 32 declined to participate after signing the consent forms, 72 failed the ‘headphone check’, and 9 others either only completed one of the *biased* sessions or did not exceed 70% accuracy on those trials that did not have distractors. We tested for this accuracy threshold during a participant’s first session, which ensured that the participants understood the basic task of categorizing three-tone sequences into two categories. All participants self-reported that they had normal hearing or hearing aids.

We had 3 study variations to randomly counterbalance the order of the session type across participants: (1) *unbiased, biased low*, and *biased high* (N=16); (2) *biased high, biased low*, and *unbiased* (N=17); and (3) *biased low, unbiased*, and *biased high* (N=15). 8 other adults (3 female; mean ± SD age of the 8 adults: 32 ± 13.95) only partially completed the study, which supplemented data in the *unbiased* experiment. Data from 5 of these participants supplemented data in the *biased low* experiments.

We collected pilot data for 9 participants and fit the full Bayesian and no-distractor models to this data. Computing the effect size based on the negative log-likelihood of the model fits, and using a power value of 0.2, we then calculated the minimum required sample size for our experiment. For this, we used the function pingouin.power-ttest (Python). The minimum required sample size was n=16.04, which indicated that our experimental sample size of 56 participants in the *unbiased* session and 48 participants across both of the *biased* sessions was sufficient to capture the effect of distractors on participant behavior. A larger cohort also helped ensure that we could study variability in the underlying factors across real-world participants. Additionally, the cohort size is similar to other comparable studies^30,41,91–93^.

### Experimental setup

During the training part of the task, participants were presented with 35 total trials. First, they were presented with 16 easy trials in which the frequencies of all three tones were signal and drawn from the mean value ± 1*σ* of either the low- or high-frequency Gaussian (signal) distributions. They were then presented with 10 more difficult trials in which two of the three tones were signal: two tones were drawn from the mean ±1*σ* of either the low- or high-frequency Gaussians, whereas the third was distractor and drawn from the extremes of the uniform distractor distribution (< *µ*_*L*_ − 4*σ* or > *µ*_*H*_ + 4*σ*). Finally, participants performed 9 very difficult trials in which only one of the three tones was signal and the other two were distractors. Before the ‘more difficult’ and ‘very difficult’ training sets, we informed the participants that the trials in those sets would comprise of one or two distractor tones respectively. Thus, the progression of the ‘easy’, ‘more difficult’ and ‘very difficult’ trials allowed us to systematically introduce the distractor tones to the participants. They were informed to ignore the distractor tones and only make a category decision based on the signal tones. We used the same training paradigm for all the 3 sessions – *unbiased, biased low*, and *biased high*. However, we did not inform them about the potentially biased nature of a session. Therefore, any information about the long-term bias was gained through exposure to the stimuli during the testing phase of the experiment.

In the testing phase, we had 600 trials in the *unbiased* session and 800 trials in each of the *biased* sessions. We randomized the trials and divided them into 4 blocks of 150 trials each for the *unbiased* session and 5 blocks of 160 trials for the *biased* sessions. The participants could optionally take a break (maximum: 3 min) between each block. Participants had to respond within 1.8 seconds after offset of the last tone and were given visual feedback about their report after each trial. The feedback duration was 0.8s, and the participants were shown either ‘Correct’, ‘Not quite’, or ‘No response. Please respond faster’. Additionally, to ensure data quality, participants were not allowed to skip or incorrectly answer 10 trials in a row; we gave them an ‘attention warning’ if 5 continuous trials were skipped or answered incorrectly. In the *unbiased* session, the median number of no-response trials was 1, and the 90^th^ percentile was 3.8. The number of no-response trials was also low during *biased low* (median: 1; 90^th^ percentile: 7) and *biased high* (median:1; 90^th^ percentile: 11.4) sessions. We excluded the no-response trials when we computed participant accuracy.

At the end of each experimental session, participants were asked for their consent to be recalled for the next session. The dates of the data collection were staggered, and we conducted each subsequent session after a variable number of days ranging from (1-30 days). We could not identify any relationship between the days between sessions and a participant’s behavior.

#### Raw Data Analysis

Fig. 2A-B: To compute the influence of each tone frequency on category choice probability 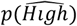, we considered the tones in the region where p(signal) > p(*distractor*). For each of these tones, we calculated the average category choice when that tone was presented as a signal tone. Then, we computed the influence as the normalized correlation coefficient between these tone frequencies and the corresponding category choice probabilities. We repeated this analysis focusing only trials where the tones were presented as a distractor tone. We additionally tested whether the position of a tone within each trial (e.g., first, second or third) affected participants’ decisions and found that these correlations were independent of tone positions (Fig S1A; Table 1, one-way repeated measures ANOVA (rmANOVA)). This analysis suggests that participants weighed all three tone positions equally within a trial and integrated information across the tones when making their category choice, and therefore our analyses could treat tone types in all three tone positions the same.

### Accuracy

We computed each participant’s accuracy by comparing their response to a given trial’s ‘true’ category. ‘True’ category is the category which is used to generate the trial using Eq. 2.

#### Computational models

##### Bayesian models

We assumed that participants performed Bayesian inference over a generative model when solving the categorization task: given their sensory evidence of the tones *m*_*d*_ where *i* ∈ {1,2,3}, participants computed the posterior probability,

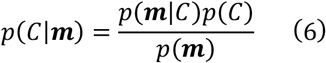

where *C* is the category choice 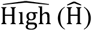 or 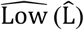. In this equation, the likelihood, *p*(***m***|*C*) that the sequence of sensory evidence ***v*** belonged to a particular category was calculated by marginalizing over a broad range of hypothesized tone frequencies ***f*** = {*f*_1_, *f*_2_, *f*_3_}. To account for the entire frequency range of human hearing^94,95^, both ***f*** and ***m*** ∈ {3.98, 39,810} Hz. In addition, we introduced a dummy variable ***r***= {*r*_1_, *r*_2_, *r*_3_} that explicitly characterized whether a participant deemed each tone’s sensory evidence (*m*_*d*_) as ‘relevant’ (denoted by R) or ‘irrelevant’ (given by I) to their category decision. Thus, the likelihood of a high-category three-tone trial is,

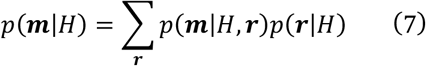

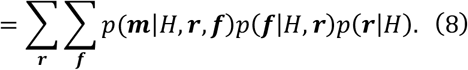

Each hypothesized tone *f*_*d*_ generated *sensory* evidence *m*_*d*_ according to the probability density *p*(*m*_*d*_|*f*_*d*_), which is a Gaussian characterized by the participant’s sensory uncertainty and noise in the auditory pathway (denoted using parameter *σ*_*sensory*_). Thus,

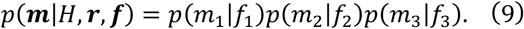

The details of how we computed the value of *σ*_*sensory*_ are given below in Methods - model fit and predictions.

Next, because the hypothesized tones and participant’s estimated relevance are conditionally independent given trial category, we can rewrite:

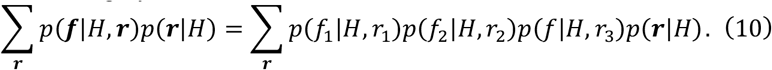

Additionally, because we observed that frequencies in each tone position are almost equally correlated with behavior (Fig S1A, S5), the relevance of a tone (denoted by parameter *p*_*distractor*_) is agnostic of its order in the three-tone sequence. Using the generative model (see Eqs. 4 and 5), we can then expand Eq. 10 as follows,

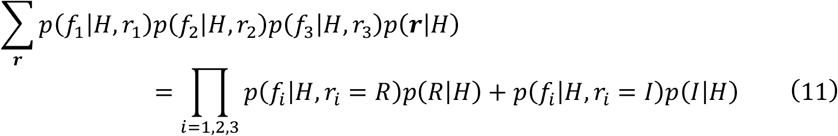

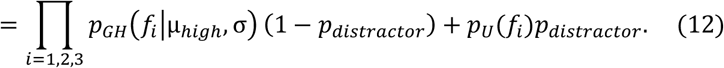

When computing the posterior in Eq. 6, *p*(*C*) = *p*_*low*_ for category choice 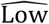 and *p*_*high*_ for category choice 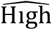. Here, *µ*_*low*_, *µ*_*high*_, *σ, p*_*distractor*_, and *p*_*low*_are the participants’ estimated values of the true experimental parameters *µ*_*L*_, *µ*_*H*_, *σ*_*expt*_, *p*_*D*_, and *p*_*L*_ and are fitting parameters in the Bayesian models. When we fit these parameters to the participants’ data, we minimized the negative likelihood (-log(posterior)), which is the cost function.

We compared three main decision-making models in the analysis: (1) A ‘full Bayesian’ model, which assumed that participants probabilistically determined if a tone was ‘signal’ or ‘distractor’; (2) a ‘no-distractor’ model, which assumed that participants considered all of the tones to be ‘signal’; and (3) a ‘random-guess’ model, which assumed that participants considered all of the tones to be ‘distractors’ and thus responded randomly. In the ‘no-distractor’ model, *p*_*distractor*_ = 0, while in the ‘random-guess’ model, *p*_*distractor*_ = 1. In these three models, stimulus relevance is agnostic of tone position (Fig 5). We also compared results from the full Bayesian model to those from an alternative model with position-dependent weights for each tone in the sequence (Fig S6).

The full Bayesian model has 6 fitting parameters – *µ*_*low*_, *µ*_*high*_, *σ, σ*_*sensory*_, *p*_*distractor*_, and *p*_*low*_– such that 0 < *p*_*distractor*_ < 1. On the other hand, in the no-distractor model, *r*_*d*_ = *R* for *i* ∈ 1,2,3 in Eq. 11. Because this model assumes that all of the tones are ‘signal’, we can simplify the likelihood *p*(***f***|*H*) into a product of three Gaussians. As a consequence, the no-distractor model has 5 fitting parameters – *µ*_*high*_, *µ*_*low*_, *σ, p*_*low*_, and *σ*_*sensory*_. In the random-guess model, participants categorized trials using random guesses proportional to the probability of the prior, and the 2 fitting parameters are *p*_*low*_and *σ*_*sensory*_. More specifically, in this model the posterior in Eq. 6 simplifies to,

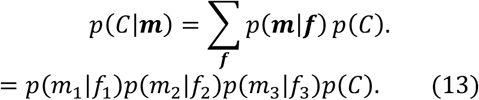

To capture change in expectations resulting from short- and long-term learning (Fig. 10), we fit an adapted full Bayesian model with a time-varying prior, such that,

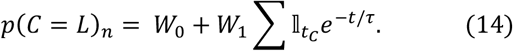

Here, *p*(*C* = *L*)_*s*_ denotes a participant’s prior for the low-category choice decision in the *n*^*dh*^ trial. The indicator function 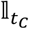 takes the value 1 if the ground truth of (*n* − *t*)^*dh*^ trial was low (L) and it takes the value -1 if it was high (H). *W*_*constant*_ = |*W*_0_ − 0.5| * 2 captures the long-term effect on participant’s expectations, *W*_1_ indicates the short-term effect and τ is the time constant governing the short-term effect.

Last, the model with position-dependent weights has 8 fitting parameters – 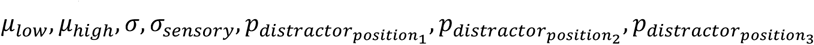 and 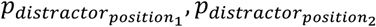 and 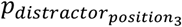 are participants’ estimates of the probabilities that the first, second and third tone respectively in the sequence are distractors.

To fit each of the above models to the participants’ psychometric curves (see Figs 6,8,9; Figs S4,5), we applied a decision threshold to the Bayesian model posterior such that,

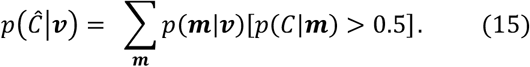

Similar to Eq. 9, we considered that each tone frequency *v*_*i*_ generated sensory evidence *m*_*i*_ according to the probability density *p*(*m*_*i*_|*v*_*i*_), which is a Gaussian characterized by the participant’s sensory uncertainty (denoted by the parameter *σ*_*sensory*_).

##### Boundary model

We constructed a simple boundary-decision model for each participant that had three model decision criteria along the frequency continuum: 1) x_L_, the boundary between low-distractor tones and low-signal tones, 2) x_C_, the category boundary between low-signal tones and high-signal tones, and 3) x_H_, the boundary between high-signal tones and high-distractor tones. Each tone was independently classified as being a distractor (*D*), low signal (*L*) or high signal (*H*). The model for individual tones can be represented as,

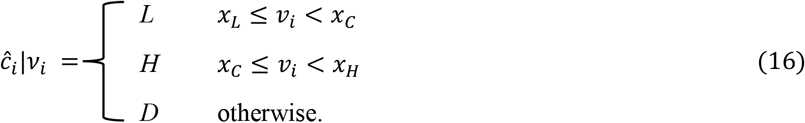

Here, ***ĉ***_*i*_ |*v*_*i*_ is the participant’s inferred generative category for tone *i*. The participant’s category report is based on the mode of the tone classifications over the three inferred tone categories ***ĉ*** (a voting model),

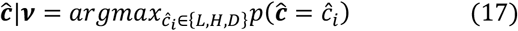

Here, c is the participant’s inferred category for the full tone sequence. The category report probability is,

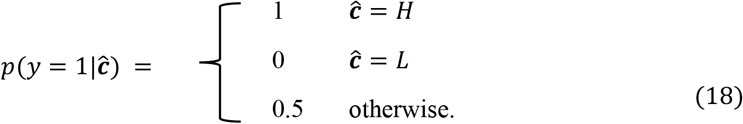

Here, y is the category report, which can be either 0 (“Low”) or 1 (“High”). If the mode of the tone classifications is “Distractor” or there is no single mode, the category report probability is 0.5.

##### Generalized linear model

We constructed an elastic net (Lasso + second-order Tikhonov) regularized binary logistic regression model^96^ for each participant such that the model predictors are the indicator functions for the three tones in a trial. An indicator function 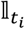 takes the value of 1 if tone *t*_*i*_ is present in the trial and 0 otherwise. There are 30 predictors because a tone frequency can have 30 possible values. The model can be represented as,

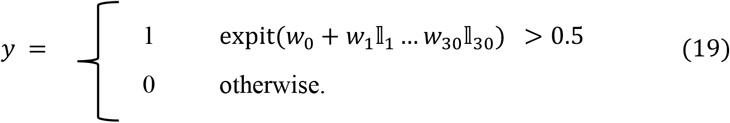

Here, *y* is the participant’s category report. The *w*s capture the influence (weight) of the different experimental tone frequencies in a participant’s decision-making process. The function expit(*x*) is computed as 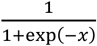.

To compute the individual influence of the three tone positions in the sequence (Fig S6H), we expand the model to have 90 predictors – 30 for each tone position.

#### Model Fit and Predictions

##### Bayesian models: full Bayesian and no-distractor models

We used two approaches to fit the full Bayesian and no-distractor models to the *unbiased* data. First, we fit all the parameters to the participant data. However, we suspected that we may be overfitting *µ*_*low*_, *µ*_*high*_, *σ* and *p*_*distractor*_, because the parameters are redundant, and there are multiple configurations that lead to the same likelihood function (Eq. 12) for the Bayesian model. Thus, in the second approach, we considered participants to be veridical about the parameters of the Gaussian distributions (*µ*_*low*_= 2.55, *µ*_*high*_ = 2.85 and *σ* = 0.1) and only fit the remaining parameters – *σ*_*sensory*_, *p*_*low*_and for the full Bayesian model, *p*_*distractor*_. Because both approaches had similar fits (see Figs S5G, H) and to prevent overfitting, we followed the second approach.

In both the approaches, we initially fit these models to a participant’s performance using a coarse parameter grid search, which allows for systematic multi-start optimization. When applicable, we used the following grid values: *µ*_*low*_∈ {2.1, 2.25, 2.4, 2.55, 2.7}, *µ*_*high*_ ∈ {2.7, 2.85, 3, 3.15, 3.3}, *σ* ∈ {0.05, 0.25, 0.45, 0.65, 0.85}, *p*_*distractor*_ ∈ {0.05, 0.25, 0.45, 0.65, 0.85} and *p*_*low*_was constrained by each participant’s correct performance. We also assumed that *σ*_*sensory*_ is participant and task dependent but not model dependent. Moreover, because the essential experimental goal was the same in both the *unbiased* and the *biased* sessions, we assumed that *σ*_*sensory*_ was constant across the three session types. Thus, instead of running a separate psychophysics experiment to compute *σ*_*sensory*_, we used data from the *unbiased* session for trials with all three tones drawn from the signal distributions and no distractor tones. We fit this data using a smaller number of model parameters – *µ*_*low*_, *µ*_*high*_, *σ, σ*_*sensory*_, and *p*_*low*_. When the remaining 4 parameters were systematically varied, we found that a participant’s *σ*_*sensory*_ was essentially constant; across the population: median – 0.19; 10^th^ percentile – 0.14; and 90^th^ percentile – 0.29. These values are consistent with previous findings^30^.

Next, like the *unbiased* session, we also fit data from the two *biased* sessions using the second fitting approach. We tested our fitting procedure by simulating 4 virtual participants (Table 2). These were modeled based on 4 actual participants from our dataset to ensure a consistency check for performance accuracy. Additionally, for the virtual participants, we computed the confidence intervals on their fitted parameters by simulating behavior 10 times and following the bootstrapping procedure as detailed below.

**Table 2:** Parameter reconstruction for 4 virtual participants.

| Participant | Parameters with fixed values |  |  | Parameters that are fitted |  |  |
| --- | --- | --- | --- | --- | --- | --- |
| | $\mu_{low}$ | $\mu_{high}$ | $\sigma$ | $\sigma_{sensory}$ | $p_{distractor}$ | $p_{low}$ |
| P1 | 2.55 | 2.85 | 0.1 | Original: 0.15<br>Fitted (median; quantile range): 0.15; 0.13 – 0.16 | Original: 0.9<br>Fitted (median; quantile range): 0.88; 0.72 – 0.97 | Original: 0.52<br>Fitted (median; quantile range): 0.52; 0.5 – 0.59 |
| P2 | 2.55 | 2.85 | 0.1 | Original: 0.23<br>Fitted (median; quantile range): 0.23; 0.2 – 0.27 | Original: 0.67<br>Fitted (median; quantile range): 0.6; 0.42 – 0.9 | Original: 0.44<br>Fitted (median; quantile range): 0.45; 0.4 – 0.49 |
| P3 | 2.55 | 2.85 | 0.1 | Original: 0.28<br>Fitted (median; quantile range): 0.28; 0.23 – 0.35 | Original: 0.88<br>Fitted (median; quantile range): 0.82; 0.46 – 0.99 | Original: 0.51<br>Fitted (median; quantile range): 0.51; 0.49 – 0.55 |
| P4 | 2.55 | 2.85 | 0.1 | Original: 0.45<br>Fitted (median; quantile range): 0.44; 0.36 – 0.53 | Original: 0.36<br>Fitted (median; quantile range): 0.35; 0.05 – 0.68 | Original: 0.46<br>Fitted (median; quantile range): 0.46; 0.43 – 0.49 |

Specifically, we generated confidence intervals by creating either 100 bootstrapped (with replacement) datasets comprising of 600 trials each for the *unbiased* session (Fig 6, Fig S5) or 100 balanced datasets for the *biased* tasks (Fig 10, Fig S10). A balanced dataset was constructed using all trials from the underrepresented category and subsampling an equal number of trials from the overrepresented category. This balanced dataset separated the external bias in the experiment (captured by *p*_*L*_ = 0.7 or *p*_*H*_ = 0.7) from the internalized expectation of the participants, as the latter can vary from 0 to 1. For example, participant in Fig 9A has negligible internalized expectation of bias, whereas participants in Figs S10C, D have high internalized expectations. These data were fit with “post-grid search”, which is a finer-resolution optimization routine using the Nelder-Mead method from scipy.optimize.minimize (Python). During this optimization, we allowed *σ*_*sensory*_ to vary within ± 0.02.

When fitting the adapted full Bayesian model to participant performance, we reduced the number of fitting parameters from 5 (*p*_*distractor*_, *σ*_*sensory*_, *W*_0_, *W*_1_ and τ) to 3 by using the median values of *p*_*distractor*_ and *σ*_*sensory*_. The parameter median values were calculated from the previous full Bayesian model fits to the bootstrapped (*unbiased*) or the balanced (*biased low* and *biased high*) datasets. To generate confidence intervals for *W*_0_, *W*_1_ and τ, for the *unbiased* data, we used 10 subsets of 500 contiguous trials and for the *biased* data, 10 subsets of 600 contiguous trials. Additionally, we fit these parameters using the following grid values: *W*_0_ ∈ {0.35,0.38 … 0.66}, *W*_1_ ∈ {0, 0.1 … 1} and τ ∈ 10^{−1,−0.85 … 0.7}^.

##### Random-guess model

Because this model has only two free parameters, *σ*_*sensory*_ and *p*_*low*_, we simplified the fitting procedure to directly fit either the 100 bootstrapped datasets (*unbiased* session) or the 100 balanced datasets (*biased* sessions). We fit *p*_*low*_with a grid search over {0, 0.1, 0.2, …, 1}.

##### Full Bayesian model with position-dependent weights

We reduce the 8 parameters in this model to 4. Participants are assumed to be veridical about the Gaussian parameters and for *σ*_*sensory*_ we use the corresponding median value. Thus, we only fit 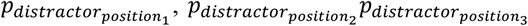 and *p*_*low*_. The fitting procedure is similar to the Full Bayesian model.

##### Metrics

Across the three experimental sessions, for nearly all the participants, we used the final model parameter values of the Full Bayesian model to compute their psychometric-curve fits and their Bayesian model posteriors (Figs 5,7-9; Figs S3,4,10). Because we had our trials had 3 tones, we plotted the average psychometric curve across all 3 tone positions.

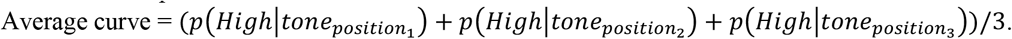

We tested the goodness of fit of the different models by comparing their Bayesian Information Criterion (BIC) values, which penalizes model complexity and encourages fits of simpler models (Figs 6; Fig S10). We also tested the results using the Akaike Information Criterion and the corrected Akaike Information Criterion^97^, which were consistent with the BIC analysis.

From the full Bayesian model fits, we derived ‘sigmoidicity’ for each participant. This metric encapsulates the nature of the shape of the psychometric curve and thus, in turn the category choice of the participant. It is ∼ 0 for participants who can accurately identify distractor tones as irrelevant to their category decisions (example participant in Fig 5B) and is ∼ 1 for participants whose psychometric curve resembles a step function. Specifically, for each participant sigmoidicity is computed using their respective posterior curve (examples in Figs S3B-D, S4B, E, H) as follows:

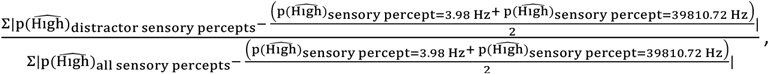

where the distractor sensory percept is more likely to be generated by the distractor tones than by the signal tones. Thus, in this formula, distractor sensory percept ∈ {3.98, μ_low_ − 2.1*σ* − median(*σ*_*sensory*_)}Hz and {μ_high_ + 2.1*σ* + median(*σ*_*sensory*_), 39810.72}Hz.

##### Generalized Linear model (GLM)

We fit a GLM to each participant’s psychophysical performance by choosing a weighting parameter α using ten-fold cross-validation. α is a parameter controlling a convex combination of Lasso and second-order Tikhonov regularization. α = 0 implies only Lasso regularization while α = 1 implies only Tikohonov regularization. For example, α for the curve fit in Fig S3E is 0.3 whereas it is 0 for those in Figs S3F and S3G. The Tikhonov matrix (τ) is designed such that τ_*i*_ = −1, τ_*i,i*−1_ = 2 and τ_*i,i*+1_ = −1.

The individual influence of the three tone positions in the sequence (Fig S5H) is computed by taking the sum of the absolute values of the corresponding GLM weights for each of the 30 tone frequencies. The weights for the first tone position are *L*_1_ … *L*_30_, for the second tone position are *L*_31_ … *L*_60_ and for the third tone position are *L*_61_ … *L*_90_. We tested our fitting procedure by simulating 6 virtual participants (Table 3). The virtual participants were based on real experimental participants.

**Table 3:** Validating the GLM for 6 virtual participants.

| Participant | $p_{\text{distractor}}$ from Bayesian model fit | Accuracy for real participants | Accuracy for simulated GLM participants |
| --- | --- | --- | --- |
| P1 | 0.58 | 74.29 | 87.17 |
| P2 | 0.68 | 85.67 | 89.5 |
| P3 | 0.7 | 82.17 | 90.5 |
| P4 | 0.79 | 83.64 | 89.67 |
| P5 | 0.89 | 82.78 | 91.33 |
| P6 | 0.99 | 82.91 | 93.17 |

##### Boundary model

We fit a boundary model to each participant’s decision behaviour by first setting the constraint *x*_*L*_ < *x*_*i*_ < *x*_*H*_. Because the stimulus set consisted of 30 unique tones, there were only 31 meaningful boundary locations. Once the order constraint was included, 4494 possible combinations of the 3 criteria remained, and the minimum negative log-likelihood was found by optimizing the model over all possible combinations.

## Data and Code Availability

The de-identified data and code associated with this paper are publicly available and can be accessed at https://github.com/geffenlab/UncertaintyRelevanceLearningInterplay. We have also used Zenodo to assign a DOI to the repository: <u>10.5281/zenodo.7439086</u>.

## Acknowledgements

We thank the members of the Geffen Lab for advice and feedback on the project. We also thank Doris Dijksterhuis and Sandra Reinert for their ideas regarding data analysis. This work was supported by the NIH grant (R01NS113241) to MNG, YC, and KK. The funders had no role in study design, data collection and analysis, decision to publish or preparation of the manuscript.

