## Supplemental Figures for "The interplay of uncertainty, relevance and learning influences auditory categorization"

**Author contributions**

J.S., J.S.C., K.P.K., E.P., Y.E.C. and M.N.G. designed the study. J.S. collected the data. J.S. and J.S.C. analyzed the data. J.S., J.S.C., K.P.K., Y.E.C. and M.N.G. wrote the paper.

**Competing interests statement**

The authors declare no competing interests in the design or execution of the study.

**Supplementary Information**

**S1 Fig.** Participants weigh tones according to their relevance; distractors adversely affect accuracy.

**S2 Fig.** Boundary model applied to 3 representative participants

**S3 Fig.** For 3 representative participants, their posteriors derived from the full Bayesian model fits and weights from a fitted generalized linear model.

**S4 Fig.** Bayesian and GLM analysis of 3 additional participants.

**S5 Fig.** Comparison of the two model fitting procedures.

**S6 Fig.** Participants' measure of stimulus relevance is tone-position dependent. However, similar to the full Bayesian model, tone relevance for all three tone positions influences category choice but not accuracy.

**S7 Fig.** Participant accuracy on the previous trial has small effect on participants' behavior.

**S8 Fig.** Participants are variable in their usage of short-term and long-term learning in the biased low and high sessions.

**S9 Fig.** The influence of long-term learning grows with time within the block.

**S10 Fig.** Most participants account for stimulus relevance in the *biased low* and *high* sessions.

**S11 Fig:** Time constant of short-term learning is inversely correlated with accuracy in the *biased low* and *high* sessions.

**S12 Fig.** Out of all the factors – sensory uncertainty, relevance uncertainty, short-term learning and long-term learning, participants' categorization accuracy is largely determined by their sensory uncertainty.

### Supporting Information

The boundary model assumes that each participant has three internal-frequency boundaries: (1) a boundary separating "low distractor" from "low signal", (2) one separating "low signal" from "high signal", and (3) a final one separating "high signal" from "high distractor" (Methods, Fig S2A). Stimuli were binned based on these internal boundaries, and we used a voting method that accounted for the three presented tones to determine the signal category.

For example, for the representative high-accuracy participant, the internal-frequency boundaries were well matched to the optimal frequency boundaries from the generative distribution (Fig S2C). However, for lower accuracy participants, the boundaries were closer together, which implies that these participants considered a wider range of the frequency space as irrelevant (Fig S2D,E).

The generalized linear model (GLM) is a descriptive model (Methods). Based on a participant's category decisions, the GLM assigns weights to each of the 30 tone frequencies (Fig S3A). The weights are indicative of the contribution of the individual tone frequencies to the participant's decision-making process. For example, for the representative high-accuracy participant, tone weights that were far from the means of the two signal categories were ~ 0 (Fig S3E, S4A), whereas those for tones that were near the category means had non-zero values. However, for participants with lower accuracy (Fig S3F,G, 4D,G), the absolute values of the weights that were far from the means of the categories were more comparable to the weights for tones near the category means.

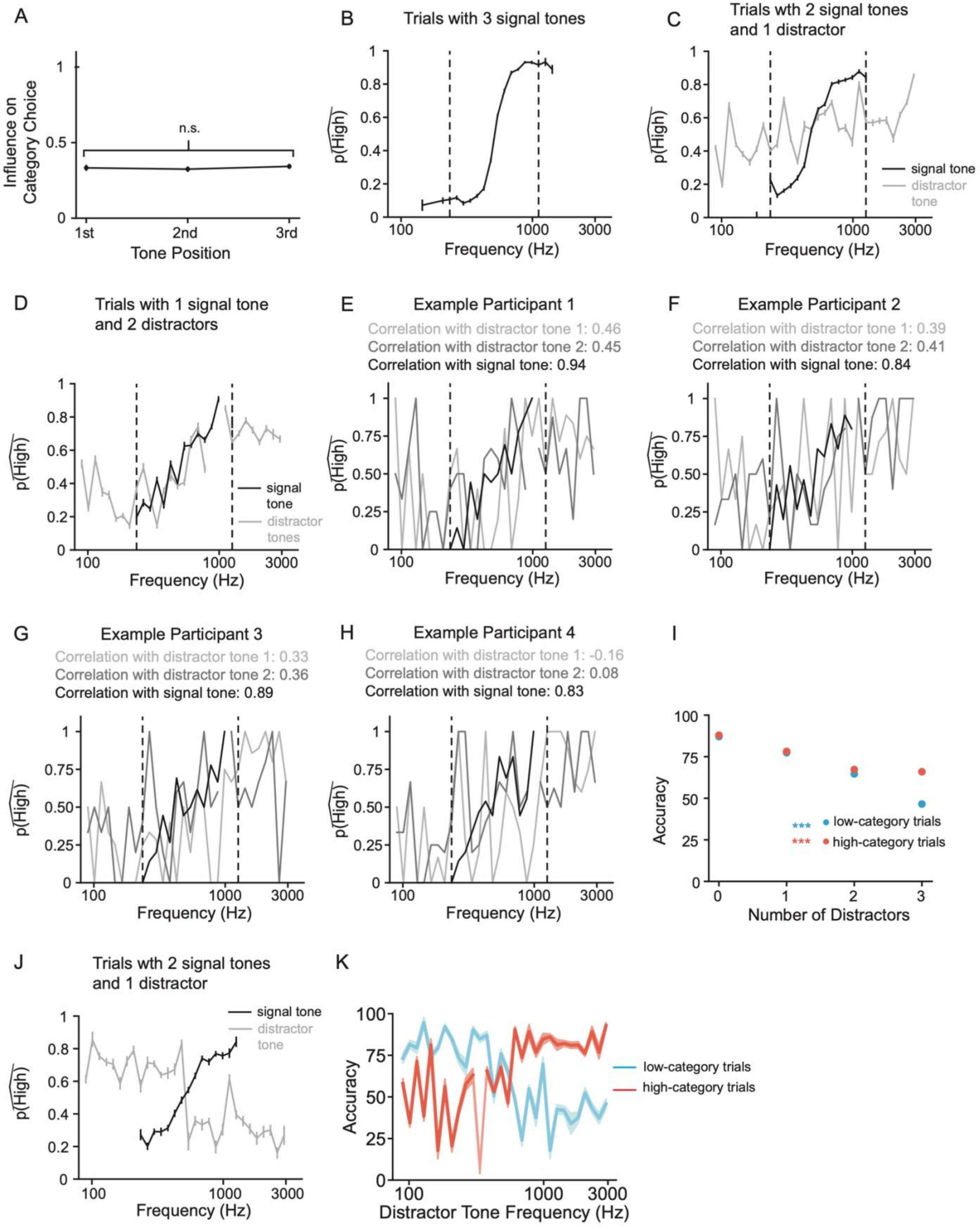
A
Influence on Category Choice
n.s.
1st
2nd
3rd
Tone Position
B
Trials with 3 signal tones
p(High)
Frequency (Hz)
C
Trials wth 2 signal tones and 1 distractor
signal tone
distractor tone
D
Trials with 1 signal tone and 2 distractors
signal tone
distractor tones
E
Example Participant 1
Correlation with distractor tone 1: 0.46
Correlation with distractor tone 2: 0.45
Correlation with signal tone: 0.94
F
Example Participant 2
Correlation with distractor tone 1: 0.39
Correlation with distractor tone 2: 0.41
Correlation with signal tone: 0.84
G
Example Participant 3
Correlation with distractor tone 1: 0.33
Correlation with distractor tone 2: 0.36
Correlation with signal tone: 0.89
H
Example Participant 4
Correlation with distractor tone 1: -0.16
Correlation with distractor tone 2: 0.08
Correlation with signal tone: 0.83
I
Accuracy
Number of Distractors
***
low-category trials
***
high-category trials
J
Trials wth 2 signal tones and 1 distractor
signal tone
distractor tone
K
Accuracy
Distractor Tone Frequency (Hz)
low-category trials
high-category trials

**S1 Fig: Participants integrate information over all 3 tones while weighing them according to their relevance; distractors adversely affect accuracy.**

(A) In *unbiased* trials, the three temporal positions had equal influence on participants' category choice i.e., the tone frequencies in each of the three positions correlated equally with participants' category choice probability. (B) Psychometric curve averaged across all participants for trials with three signal tones. Because a given signal tone frequency can be played in any order (as the 1st, 2nd or 3rd tone) in a trial sequence, here, we plot the mean of the three resulting $p(\widehat{High})$ values. Average psychometric curves plotted separately for signal and distractor tones (C) for trials with one distractor tone and (D) for trials with two distractor tones. Also, to compute the correlations in Figs 2 and 3, we only consider the frequency range – marked using black dashed lines – where the category Gaussian probability distributions are greater than the distractor uniform distribution. This allows us to compare signal and distractor correlations across the different trial types. (E-H) Correlations for trials with two distractor tones for 4 example participants. (I) Accuracy for trials from both categories is inversely correlated with number of distractors (Statistics: one-way repeated measured ANOVA, $p_{tone\_position\_low\_category} = 3.03e\text{-}62$, $F_{tone\_position\_low\_category}(3, 165) = 259.64$, $p_{tone\_position\_high\_category} = 7.71e\text{-}29$, $F_{tone\_position\_high\_category}(3, 165) = 68.56$; Table 1). (J) Average psychometric curve for signal and distractor tones when the distractor tone frequency is incongruent with the trial category, plotted for trials with only one distractor. (K) Similar to 2C, for trials with 2 distractor tones, accuracy was higher when the distractor tone frequencies were similar to the trial category. In panels (B-D, I-K), mean and SEM over N=56 participants; ***$p$<0.0001.

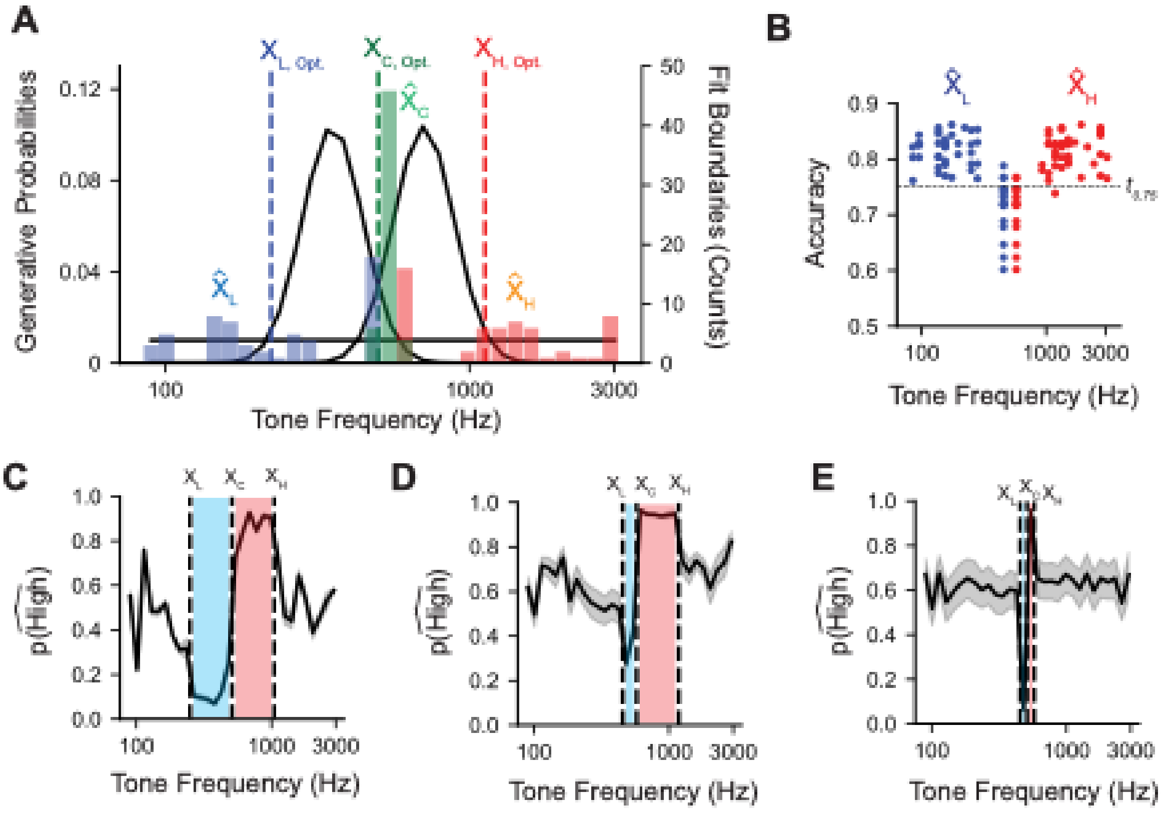


**S2 Fig: Boundary Model applied to three representative participants.**
(A) The optimal boundary locations (vertical dashed lines; $X_{L, Opt}$, $X_{C, Opt}$, and $X_{H, Opt}$,) delineate the low, central, and high frequency regions, respectively, in which generated tones are most likely to be from a specific distribution. We fit these boundaries to individual participants and plotted the distributions of the best-fitting low, center, and high boundaries ($\widehat{X_L}$, $\widehat{X_C}$, and $\widehat{X_H}$,). The best-fitting center boundaries are centered on the optimal boundary location, whereas the high and low boundaries are skewed towards the extremes, indicating an under-estimation of $p_{distractor}$. (B) A subset of (N = 10/56) were best fit by low and high boundaries adjacent to the center boundary. These "poor fit" participants perceived most of the stimulus space as distractors. We removed poor-fit participants from this plot. (C-E) Simulated data based on the fits to representative participants in Figure 5. Error bars: SEM. Blue shaded area indicates the frequency range that is considered to be low signal, whereas the red shaded area indicates the frequency range that is considered to be high signal.

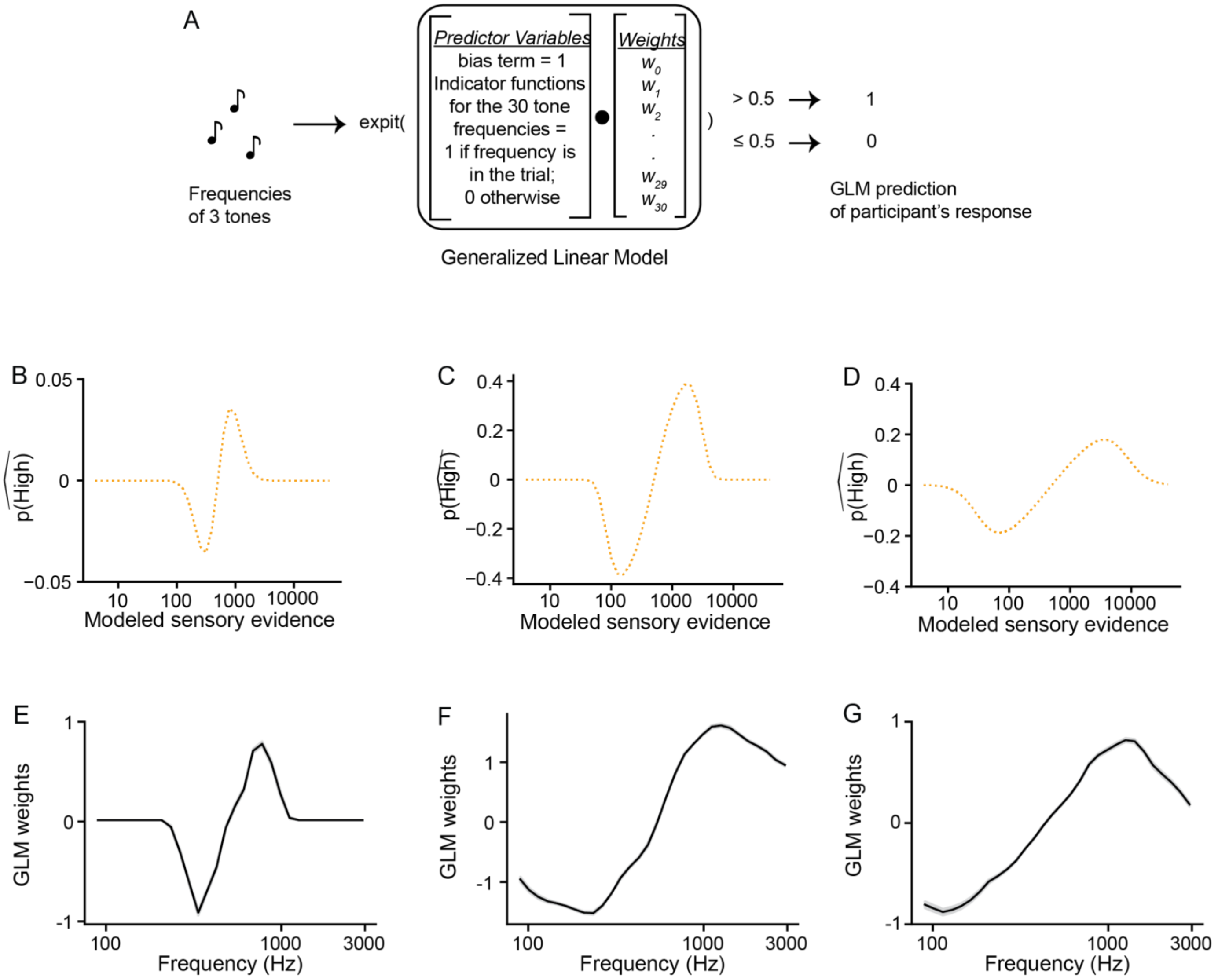


**S3 Fig: For 3 representative participants, their posteriors derived from the full Bayesian model fits and weights from a fitted generalized linear model.**

(A) Schematic of the generalized linear model (GLM) that has the frequencies of the three tones in a trial as input and uses the corresponding indicator functions as its predictors to predict the participant's category choice response (see Methods). (B-D) Posteriors generated from the full Bayesian model fits to the representative participants' data in Fig 5. The x-axis is a participant's sensory evidence i.e., their internal representation of the different tone frequencies. The average posterior for each participant is generated in a manner similar to their average psychometric curve. Error bars: standard error of the mean; not large enough to be distinguishable. (E-G) GLM weights of tone frequencies in the respective participant's category decision. Mean and error bars of the curve: computed using cross-validation (n=10, see Methods).

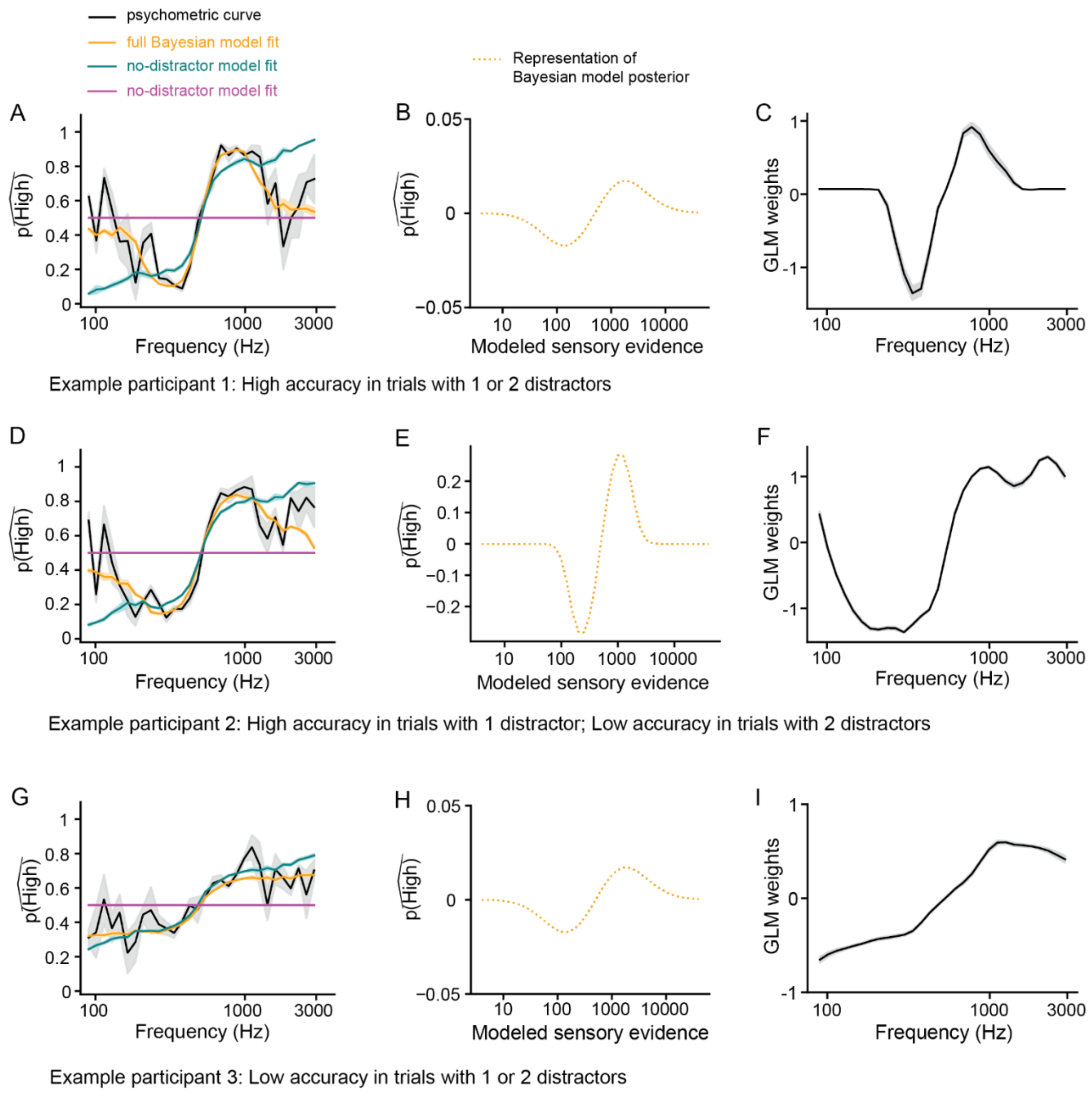


**S4 Fig: Bayesian and GLM analysis of 3 additional participants.**

(A, D, G) Similar to Fig. 5 psychometric curves for 3 additional participants; black lines: raw data, orange lines: full Bayesian model fit, teal lines: no-distractor model fit, magenta lines: random-guess model fit. (B, E, H) Posteriors generated from the full Bayesian model fits to performance in (A, D, G) respectively. (C, F, I) Relative weights of the tone frequencies determined using the GLM. For all subplots the average curve and its error bars are obtained as described in the captions for Fig 5 and Fig S3. (A, B, C): example participant with high task accuracy in trials with either one or two distractors; one distractor: 86.55, two distractors: 72.17. (D, E, F): example participant with high task accuracy in trials with one distractor, and poor accuracy in trials with two distractors; one distractor: 82.91, two distractors: 64.91. (G, H, I): example participant with poor accuracy in both trials with one or two distractors; one distractor: 57.63, two distractors: 64.66.

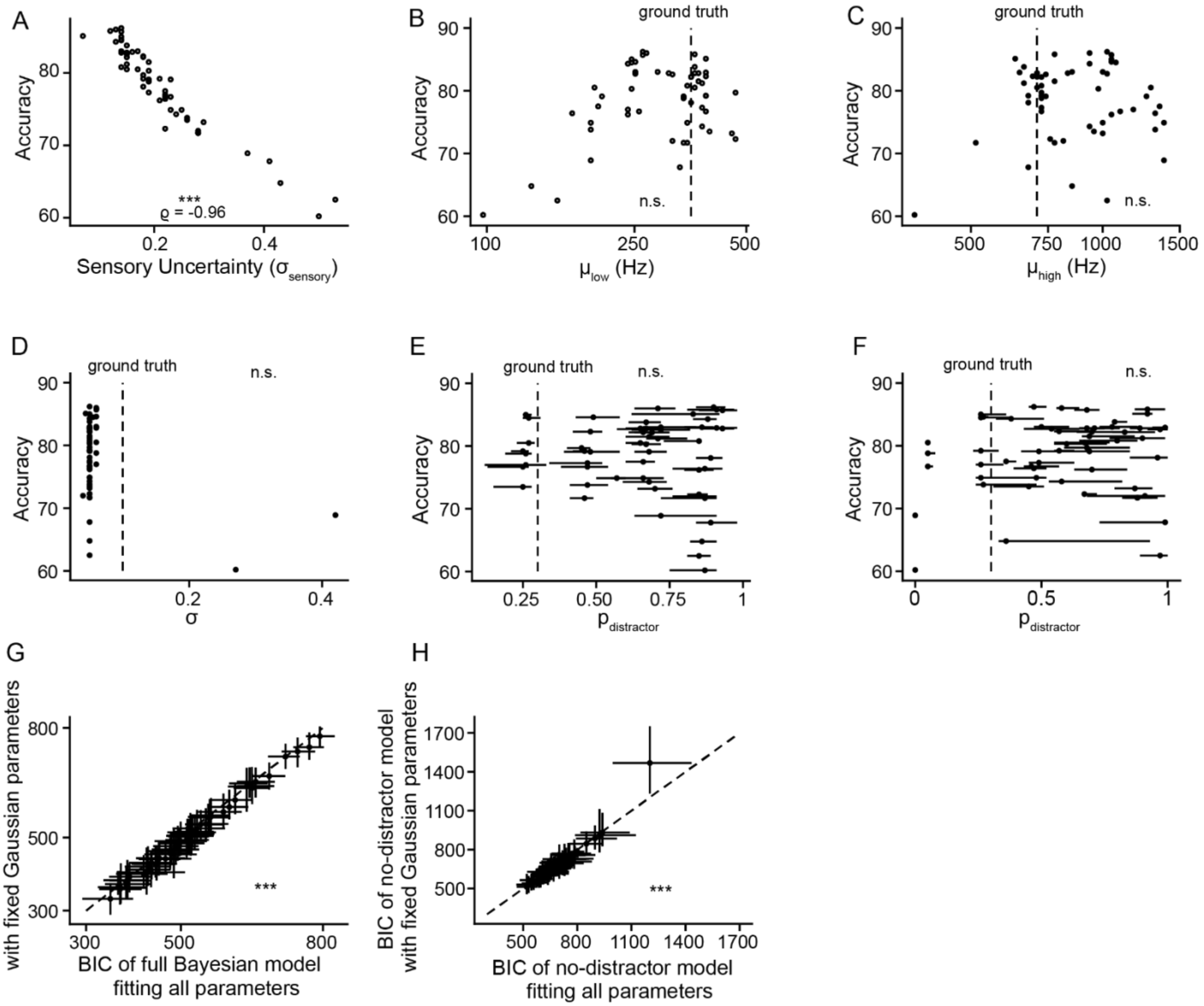


**S5 Fig: Comparison of the two model fitting procedures.**

(A) Values for $\sigma_{sensory}$ (i.e., sensory uncertainty) are the same for both fitting procedures and are inversely correlated with accuracy. (B-E) Parameter values obtained from fitting all 6 parameters of the full Bayesian model to participants' data. Parameters (B) $\mu_{low}$, (C) $\mu_{high}$, (D) $\sigma$ of the two Gaussians, and (E) $p_{distractor}$ are uncorrelated with accuracy. (F) corresponds to the parameter values of $p_{distractor}$ obtained using the second fitting procedure wherein participants are considered veridical about the Gaussian parameters. Vertical lines in (B-E) denote 'ground-truth' values, which are the true parameter values used in the generative process. Panels (A-F) Statistics: Spearman Correlation (Table 1). (G, H) Comparison of the two fitting approaches: for most participants BIC values of the full Bayesian and no-distractor models from the two approaches are roughly similar although fitting all parameters results in slightly higher BIC values. (full Bayesian model, fitting all parameters: Mdn: 514.99, IQR: 115.13, fixing Gaussian Parameters: Mdn: 504.21, IQR: 127.23; no-distractor model, fitting all parameters: Mdn: 660.13, IQR: 111, fixing Gaussian Parameters: Mdn: 659.66, IQR: 117.39) Statistics: two-tailed Wilcoxon signed-rank test comparing BIC scores from the two fitting procedures (Table 1). In all panels, data point: one participant; $^{\dagger}p<0.05$; $^{*}p<0.01$, $^{**}p<0.001$, $^{***}p<0.0001$.

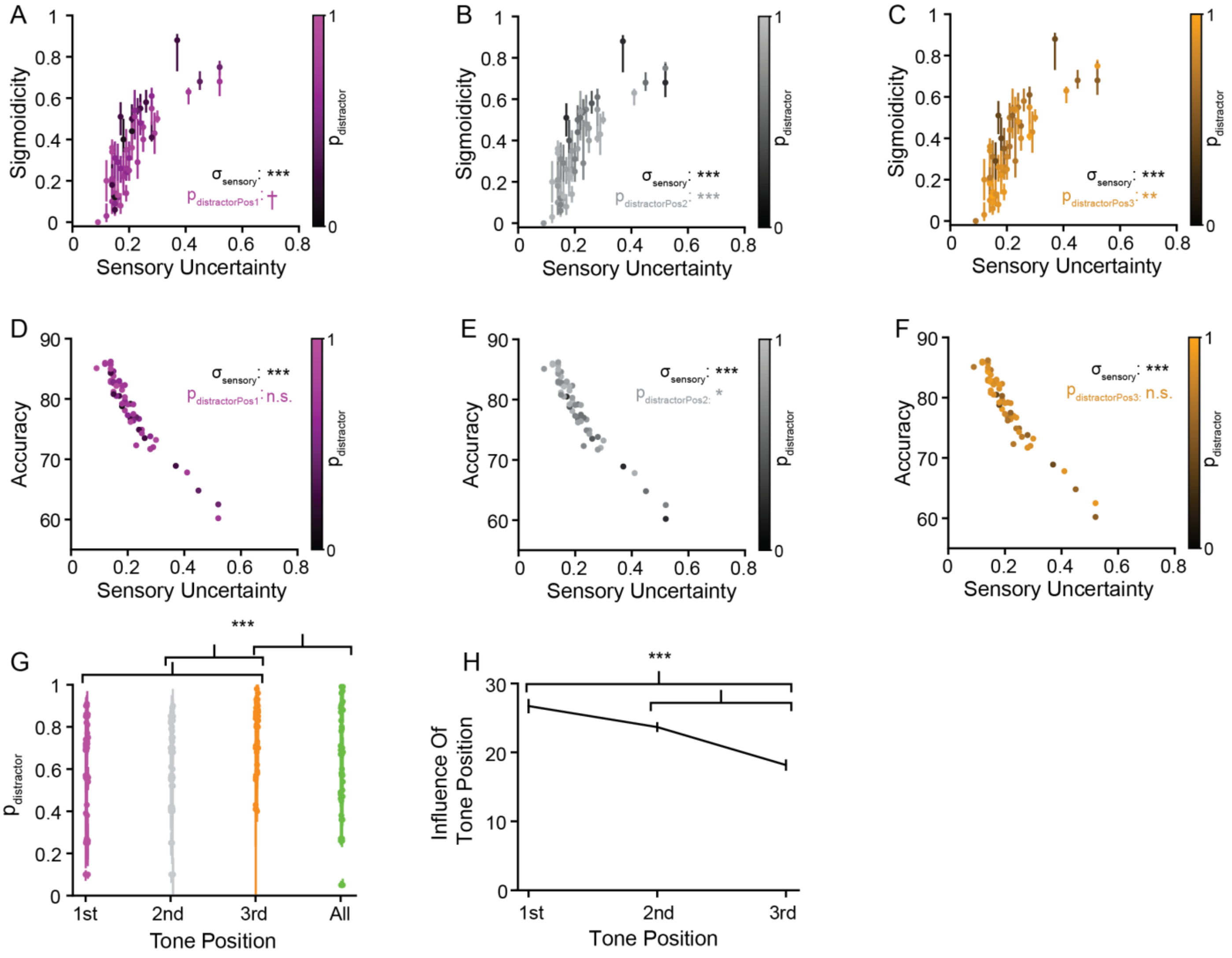


**S6 Fig: Participants' measure of stimulus relevance is tone-position dependent. However, similar to the full Bayesian model, tone relevance for all three tone positions influences category choice but not accuracy.**

(A-C) Similar to Fig. 6B, both $p_{distractor}$ and $\sigma_{sensory}$ affect 'sigmoidicity'; the different colors correspond to the three tone positions as denoted in (G). Statistics: Spearman correlation of respective parameter with 'sigmoidicity'. (D-F) Similar to Fig. 6C, $\sigma_{sensory}$ but not the position-dependent $p_{distractor}$ values influence accuracy; the colors are consistent with (G). Statistics: Spearman correlation of respective parameter with accuracy. (G) Participants' estimates of the probability of distractors are statistically similar for the first ($p_{distractor_{position_1}}$) and second ($p_{distractor_{position_2}}$) tone positions but are higher for the third ($p_{distractor_{position_3}}$) tone position (one-way rmANOVA; Statistics: two-sided Wilcoxon signed-rank test). Only estimates for the third position are different from those ($p_{distractor}$) derived using the full Bayesian model. (H) Influence of the three tone positions computed using the Generalized Linear Model (see Methods). Statistics: two-sided Wilcoxon signed-rank test. In panels (A-G), data point: one participant; †p<0.05; *p<0.01, **p<0.001, ***p<0.0001.

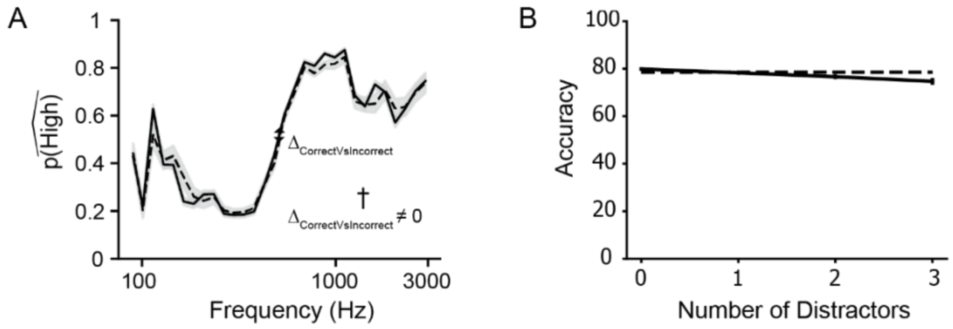


**S7 Fig: Participant accuracy and number of distractors on the previous trial have small effect on participants' behavior.** (A) Average psychometric curves conditioned on whether previous choice was correct versus incorrect. Dashed line: previous trial is incorrect, solid line: previous trial is correct. $\Delta_{\mathrm{CorrectVsIncorrect}}$: ~0.019; $\Delta_{\mathrm{CorrectVsIncorrect}}$: difference between the two psychometric curves at the central frequency. Error bars: standard error of the mean. Statistics: two-tailed Wilcoxon signed-rank test on $\Delta_{\mathrm{CorrectVsIncorrect}}$ (Table 1), †p<0.05. (B) Accuracy decreases slightly with increase in number of distractors in the previous trial. However, these accuracies are approximately equal to the overall accuracy of 78.4% (dotted black line). Error bars: standard error of the mean. Statistics: one-way repeated measured ANOVA, *p = 3e-6*, $F_{tone\ position}(3, 165) = 10.35$. (Table 1)

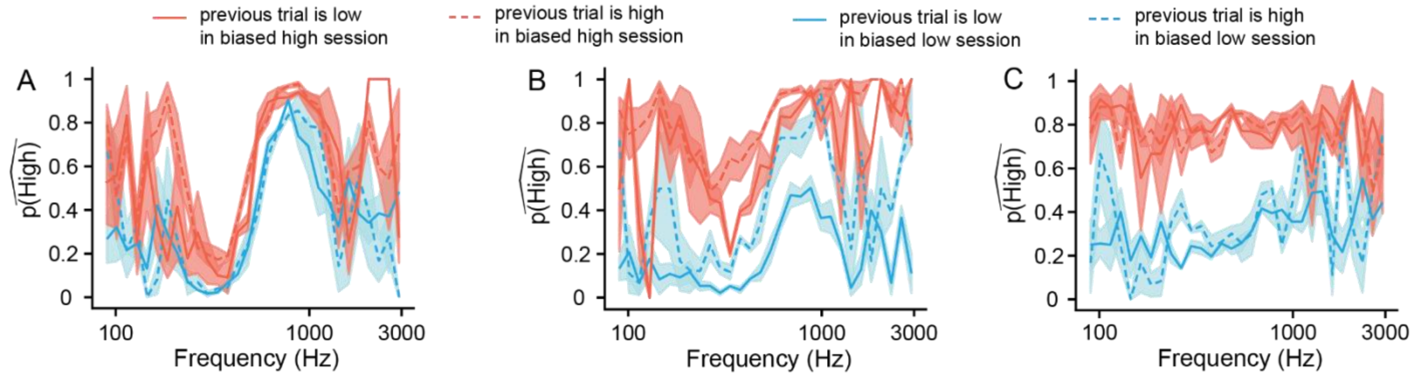


**S8 Fig: Participants are variable in their usage of short-term and long-term learning in the *biased low* and *high* sessions.** (A-C) Similar to Figs. 8A-C, average psychometric curves conditioned on the previous trial's category type for three participants in the *biased low* (light blue) and *biased high* (red) sessions. These curves are from the same participants as Figs. 9B-D. (A) Example participant who anecdotally does not demonstrate any reliance on the previous trial category nor on session-level bias. (B) Example participant who relies on both the previous trial category and on session-level bias. (C) Example participant who relies on session-level bias but not on the previous trial category. $\Delta_{short-term}$: (A) *biased high* session: 0.04, *biased low* session: -0.02. (B) *biased high* session: 0.19, *biased low* session: 0.28. (C) *biased high* session: -0.01, *biased low* session: 0.02.

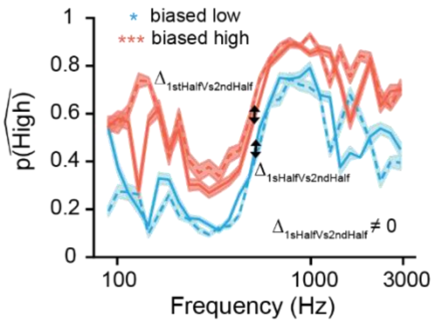


**S9 Fig: The influence of long-term learning grows with time within the block.**
Psychometric curves for the *biased low* (light blue) and *biased high* (red) sessions, averaged across all participants and conditioned on time within the block; $\Delta_{1stHalfVs2ndHalf}$: ~0.033 (for *biased low*) and $\sim -0.063$ (for *biased high*). $\Delta_{1stHalfVs2ndHalf}$: difference between the two psychometric curves for each respective session at the central frequency. Statistics: two-tailed Wilcoxon signed-rank test on $\Delta_{1stHalfVs2ndHalf}$ (Table 1), $^{\dagger}p<0.05$; *$p<0.01$, **$p<0.001$, ***$p<0.0001$. Error bars: standard error of the mean.

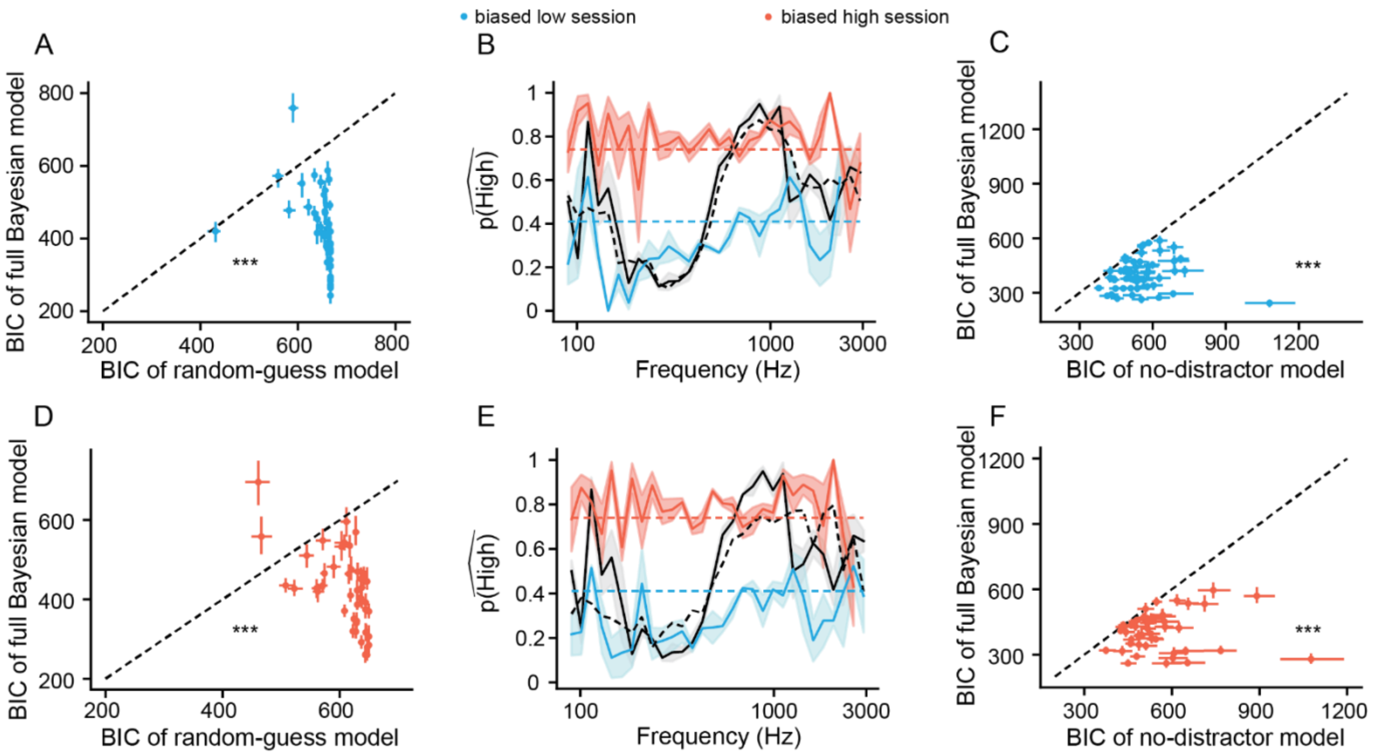


**S10 Fig: Most participants account for stimulus relevance in the *biased low* and *high* sessions.**

(A) BIC scores of the full Bayesian model are lower than those of the random-guess model for all except 2 participants in the *biased low* session (light blue dots). Data point: one participant; number of participants = 53. (D) Corresponding plot for the *biased high* session (red dots); number of participants = 48. In (A, D) Error bars: 95% confidence interval. Average BIC scores and error bars are obtained using model fits to 100 balanced datasets (see Methods; Table 1). Statistics: two-sided Wilcoxon signed-rank test comparing the two model fits (Table 1). (B, E) Psychometric curves for the two participants that have a higher BIC value for the full Bayesian model compared to the random-guess model. Average curves for *biased* sessions (light blue curve – *biased low*, red curve – *biased high*) are evaluated using 20 subsampled balanced datasets. Error bars: standard error of the mean. Curves for the *unbiased* session, shown using black lines, are similar to Figs. 5B-E. Respective full Bayesian and random-guess model fits are shown using dotted lines. (C, F) For all other participants besides these two, BIC scores of the full Bayesian model are lower than those of the no-distractor model for both the *biased* sessions. Statistics: two-sided Wilcoxon signed-rank test comparing the two model fits (Table 1). In all plots: ***$p<0.0001$.

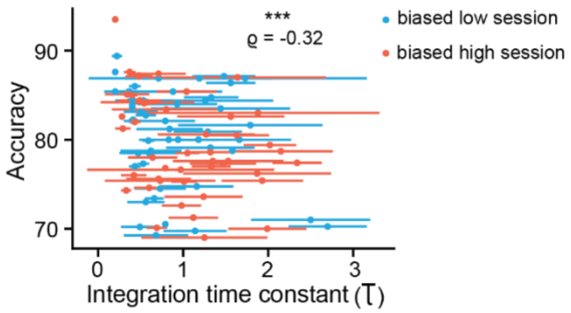


**S11 Fig: Time constant of short-term learning is inversely correlated with accuracy in the *biased low* and *high* sessions.**
For participants whose performance was fit using the adapted full Bayesian model (biased low: light blue, N=51; biased high: red, N=46), similar to the *unbiased* trials (Fig 10B), their short-term learning time constant was inversely correlated with their accuracy. Statistics: Spearman correlation; *p< 0.01

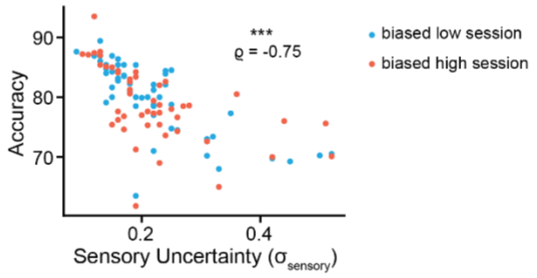


**S12 Fig: Out of all the factors – sensory uncertainty, relevance uncertainty, short-term learning and long-term learning, participants' categorization accuracy is largely determined by their sensory uncertainty.**
For all participants, similar to Fig. 7C, category accuracy was impaired by sensory uncertainty. Data points: one participant; Error bars: standard error of the mean; Statistics: Spearman correlation; ***$p<0.0001$
